# Warming amplifies short-term responses to disturbance in lake food webs

**DOI:** 10.64898/2026.07.21.739822

**Authors:** G. Virginio Clemente, Corey J. A. Bradshaw, Giovanni Strona

## Abstract

Ecological communities can exhibit transient amplification after disturbance even if asymptotically stable, yet how environmental change reshapes these short-term responses in real food webs remains unclear. We combined four decades of monthly plankton observations from nine Swiss lakes with a food-web model to quantify effects of warming and oligotrophication on two transient properties: reactivity (instantaneous sensitivity to small disturbances) and the time to peak response (*t*_max_). Across lakes, community states lie in an asymptotically stable yet reactive regime. Warmer conditions consistently increased reactivity and lengthened *t*_max_; after accounting for oligotrophication, a 1 °C rise in mean temperature corresponded to ∼ 10% higher reactivity on average. Linking month-to-month variation in transient metrics to guild biomass showed that reactivity covaried strongly with consumer biomass, especially large herbivores, whereas *t*_max_ was most associated with predator biomass, a pattern consistent with metabolic-scaling predictions from thermal ecology. The central warming-reactivity association was preserved when we made the bioenergetic parameters themselves temperature-dependent through Boltzmann-Arrhenius scaling. Warming therefore systematically boosted transient sensitivity in these lake food webs via shifts in consumer and predator biomass.

**Author summary:** Ecological stability is often evaluated using long-term equilibrium properties that describe whether a community eventually returns toward its previous state after a disturbance, and how long this recovery takes. Yet a system that is stable in the long run can still undergo a strong short-term response before recovering. We asked whether environmental warming changes the magnitude and duration of these temporary responses in real ecosystems. We combined four decades of monthly plankton observations from nine Swiss lakes with a mathematical model describing interactions among major feeding groups. We found that warmer conditions were consistently associated with stronger responses to disturbance and with a longer delay before those responses reached their maximum. These patterns emerged despite the communities remining stable over the long term. Changes in the amount of biomass held by consumers and predators helped explain the observed responses, suggesting that warming alters stability through changes in food-web structure. Our findings show that long-term measures alone may overlook ecologically important short-term dynamics. Considering both eventual recovery and temporary amplification could therefore improve our understanding of how lakes and other ecosystems respond to ongoing climate change and increasingly frequent disturbances.

## Introduction

Climate change is reshaping ecological communities by altering the temperature dependence of organismal physiology, including metabolic and ingestion rates [1, 2], with consequences for ecosystem metabolism and trophic energy transfer [3, 4]. At the same time, climate forcing modifies the abiotic context in which communities are embedded, including nutrient regimes, mixing, stratification, hydrology and light availability [5–7]. Food webs, directed networks of trophic interactions, represent the natural backbone of these communities and provide a powerful framework to describe how ecological interactions and energy pathways reorganize under sustained environmental forcing [8–10]. An open question is how sustained warming changes the way real food webs respond to disturbance, especially on the short timescales over which climate-driven perturbations such as heat waves, stratification anomalies, and runoff pulses are increasingly likely to occur [6, 11, 12].

Theory and microcosm experiments have established that temperature influences several ingredients of community dynamics simultaneously, but unequally. Respiratory and consumer metabolic demands often increase faster with temperature than photosynthesis or primary production [3, 4, 13], and ingestion rates, attack rates, and handling times also respond to temperature, but with rate- and taxon-specific sensitivities [14, 15]. When warming increases consumer energetic demand faster than resource production, predicted consequences include stronger top-down control, intensified per-capita interaction strengths, and under some model assumptions, destabilization of long-term equilibria [16–18]. However, these outcomes are not universal: food-web structure, predator interference, and functional-response shape can dampen enrichment- or warming-driven instability [19]. Empirical work has begun to confirm parts of this picture: warming-induced metabolic plasticity can amplify ecosystem-level energy fluxes [2], and sustained climate stress can reorganize trophic pyramids when food-web architecture fails to compensate [9].

Most of this work has focused on long-term or asymptotic measures of stability, commonly summarized for linearized dynamics by the dominant eigenvalue *α* of the Jacobian, the matrix describing how small changes in the biomass of each trophic guild instantaneously affect the rates of change of the others. The sign of *α* indicates whether sufficiently small perturbations ultimately decay or grow [20–22]. But for systems exposed to repeated, climate-driven perturbations, the more consequential dynamics may unfold on much shorter timescales [23–25]. Even when an equilibrium is asymptotically stable, perturbations around it can be transiently amplified, growing in magnitude before eventually decaying [26–29]. This transient amplification is captured by the *reactivity* (*ρ*), the maximum instantaneous growth rate of a small perturbation, while its timing is described by the time at which the amplification peaks (*t*_max_). These complementary quantities are illustrated in Fig. 1.

**Fig 1.**
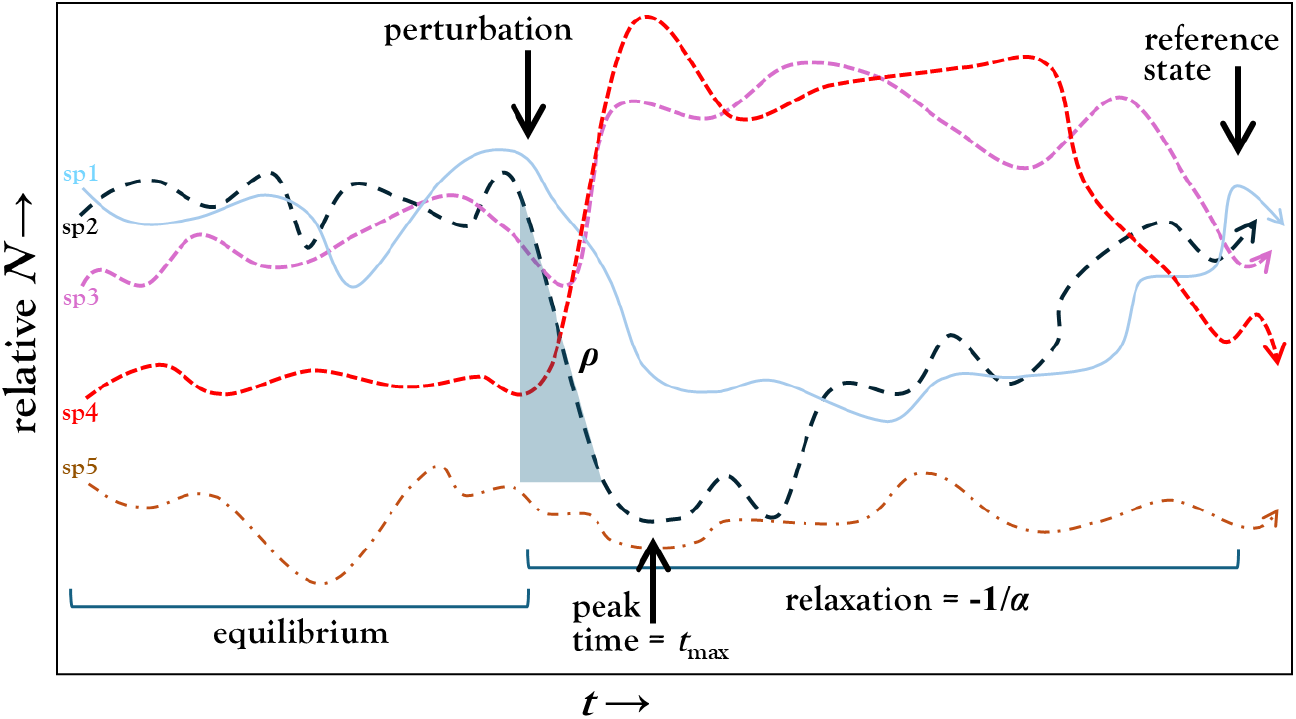
Transient dynamics following a small perturbation around an equilibrium in a reactive system. The time series show the temporal evolution of five species’ relative abundances (*N*) after perturbation. The main features of the transient response are indicated: reactivity (*ρ*), peak time (*t*_max_), and relaxation time (−1*/α*) toward the reference state, where *α* denotes asymptotic stability.

Plankton communities offer a clear setting for studying such short-term responses: disturbance can produce rapid biomass accumulation during blooms [30] or abrupt crashes in major plankton compartments [31], followed in some systems by recovery toward recurrent or predisturbance community states [6, 32]. Even temporary excursions from equilibrium can reshape trophic control and affect persistence, making transient dynamics important for the estimation of extinction risk [24, 29].

Despite this growing theoretical interest, the empirical picture remains incomplete. Most work on reactivity has relied on analytical theory, random-matrix models, or simulated networks [27, 29], but long-term, multispecies field tests are rare. Moreover, *ρ* and *t*_max_ are rarely considered jointly, although they answer complementary questions: how much perturbations are amplified and how long amplification lasts. It is also unclear which trophic guilds drive observed changes in transient dynamics, and whether warming-related changes in these metrics follow the consumer-mediated pathways predicted by metabolic theory.

Metabolic theory provides specific expectations for these pathways. Warming alters consumer and predator metabolic demand and feeding-related rates, often differently from basal resource production [1, 13, 14, 16, 33]. These asymmetric responses can modify consumer-resource interaction strengths and top-down control; where they strengthen interaction channels that transiently amplify perturbations, reactivity is expected to increase [27, 34]. Temperature can also alter generation times and predator-prey response times asymmetrically [15, 33], suggesting that the timing of peak amplification, *t*_max_, may shift with warming, although theory alone does not predict its direction. Finally, because transient amplification depends on the strength and organization of interactions in the Jacobian, variation in *ρ* and *t*_max_ should be expressed particularly through consumer and predator guilds rather than through primary producers alone. We test these predictions using four decades of monthly plankton observations across nine Swiss lakes [35], combined with a bioenergetic food-web model [36] that lets us evaluate the local Jacobian at each monthly community state. The dataset comprises time series for trophic guild biomasses linked by literature-derived predator-prey relationships (Fig. 2). The lakes have experienced two contrasting environmental trends since the mid-twentieth century: temperatures have risen steadily since the 1950s, while re-oligotrophication policies initiated in the 1970s have progressively lowered dissolved phosphate concentrations. The magnitudes of these changes differ across lakes (Fig. 3), creating a natural contrast in how warming and nutrient decline jointly shape food-web dynamics.

**Fig 2.**
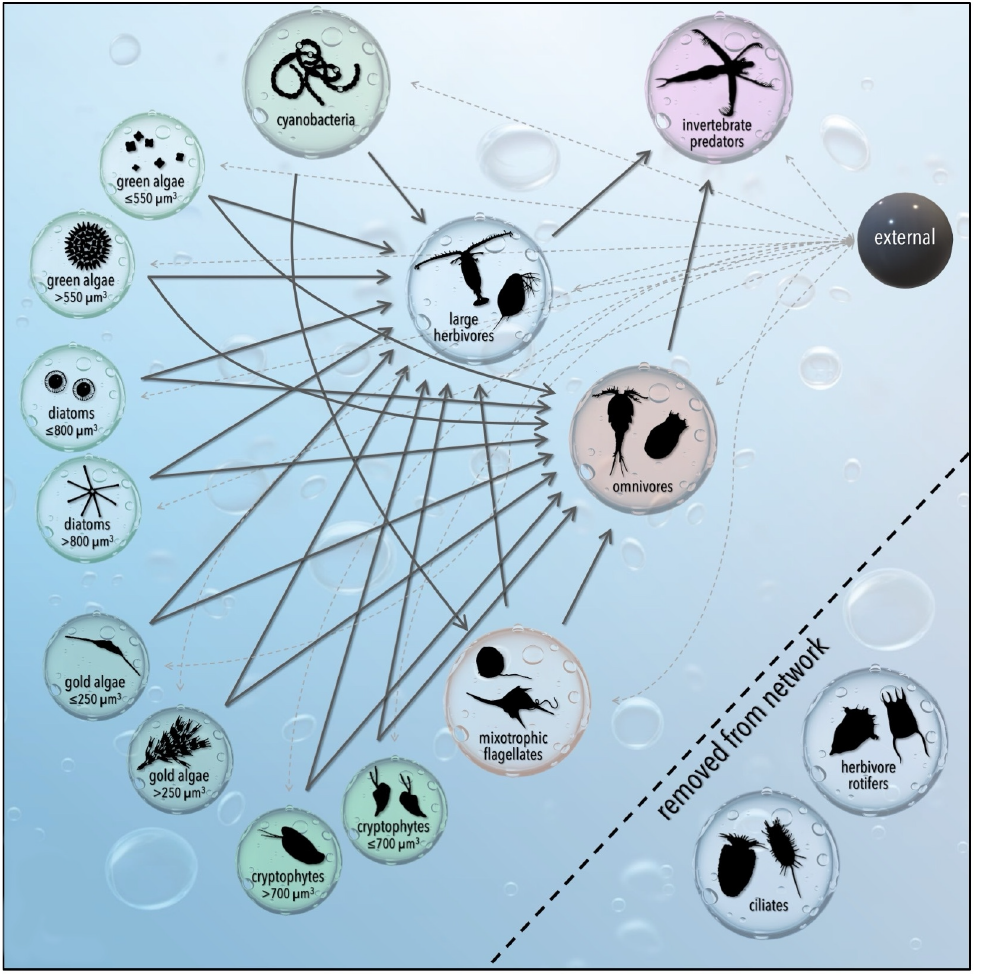
Conceptual representation of the planktonic food web. The diagram shows trophic groups and directed predator-prey interactions, including links to an external node. Nodes excluded from the analyzed network are indicated.

**Fig 3.**
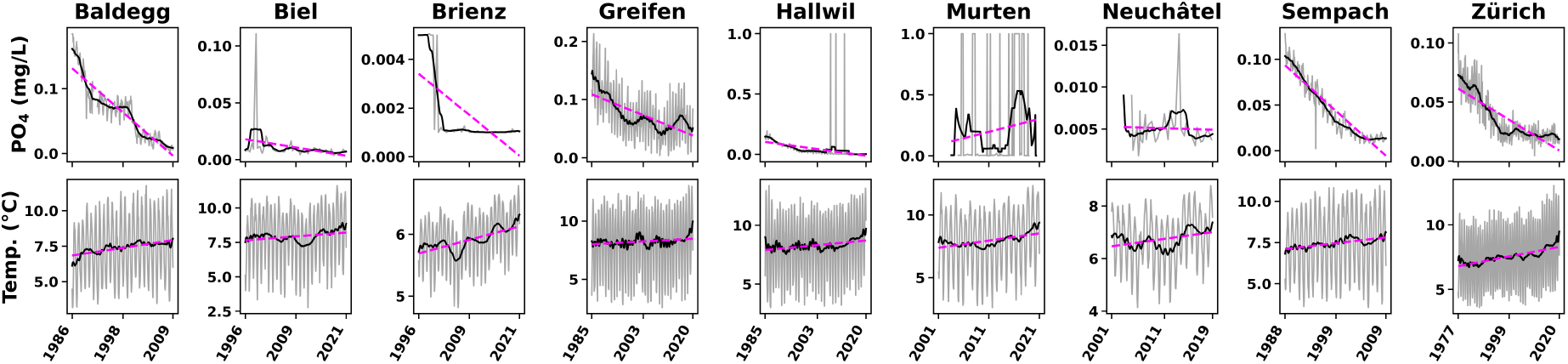
Temporal trends in phosphate concentration and water temperature across the studied lakes. Top: phosphate concentration (PO_4_, mg/L). Bottom: water temperature (°C). Grey lines represent monthly observations, solid black lines show 36-month rolling averages, and dashed magenta lines indicate linear trends over the study period.

Using published metabolic and growth parameters [36], we reconstructed local community dynamics and quantified two complementary properties of transient responses: reactivity (*ρ*) and the time to peak response (*t*_max_). Rather than assuming long-term equilibrium, we treated each monthly observation as a state-dependent operating point of a continuously forced system and computed transient metrics from the Jacobian evaluated at that state. Throughout the period for which there are data, inferred states sat in a regime that was asymptotically stable yet reactive (S1 Fig): perturbations decay in the long run, while transient amplification remains possible.

## Materials and methods

### Modelling framework: from biomass time series to transient metrics

Our analysis combines two conceptually distinct modelling steps. We first used a bioenergetic food-web model [36] to map each monthly community state to a set of transient stability metrics, and then build time-series models to ask how those metrics covary with environmental drivers. Because the interplay between these two steps is central to interpreting the results, we summarize it here before detailing each component.

For each lake-month, we combined observed biomasses ***B***(*t*) with the bioenergetic model to construct the local Jacobian ***J*** (*t*) of the linearized dynamics; from ***J*** (*t*) we extract three scalar descriptors of the response to small perturbations: asymptotic stability (*α*(*t*)), reactivity (*ρ*(*t*)), and the time to peak response (*t*_max_(*t*)). These are model-derived quantities, not directly observed. We chose the Boit et al. [36] parameterization because it was developed and validated specifically on a similar plankton food web (Lake Constance) using empirical biomass data, which makes it the most defensible parameterization for our lakes. We did not integrate the equations forward in time; we used them only to define the local linearization and the implied transient response properties.

We take the time series *ρ*(*t*) and *t*_max_(*t*) as response variables in seasonal autoregressive integrated moving-average models with exogenous regressors (SARIMAX) [37], with monthly lake temperature and phosphate (and their lags) as exogenous predictors. This time-series step is appropriate here because both response and predictor series are strongly autocorrelated and seasonal; ignoring this structure would inflate apparent associations. The approach establishes whether month-to-month variation in environmental drivers is systematically associated with variation in non-equilibrium stability metrics, after accounting for temporal structure. It cannot, on its own, propagate the parameter uncertainty embedded in the bioenergetic model into the SARIMAX coefficients; we therefore interpret SARIMAX coefficients to describe how warming relates to model-implied transient metrics, and we provide a sensitivity analysis (below and S1 Appendix, Section SI-G) in which we made the bioenergetic parameters explicitly temperature-dependent.

### Data

We extracted plankton abundance data from Merz et al. [35] for nine Swiss lakes (S8 Fig). The dataset comprises monthly biomass time series spanning 20-40 years for fifteen functional groups (including large invertebrate predators, omnivores, large herbivore grazers, small grazers, mixotrophs, and several primary producers guilds). We excluded rotifers and ciliates from the analysis because these small-grazer groups were not consistently quantified across lakes; Merz et al. [35] showed that connectance and interaction-strength patterns were similar with and without these groups. Trophic topology followed the guild-level scheme of Boit et al. [36], restricted to prey-predator links (Fig. 2). Each time series is paired with monthly records of water temperature and dissolved phosphate; for more details see [35].

Plankton communities are continuously perturbed by storm-mixing events, heat-wave-induced stratification anomalies, sudden nutrient pulses, pulses of organic matter, and stochastic recruitment failures [6, 30–32]. We did not seek to identify or quantify these perturbation events; rather, we asked what the local food-web configuration at each monthly observation implies for the system’s susceptibility to a small perturbation of arbitrary direction. The transient metrics *ρ* and *t*_max_ are properties of the linearized operator ***J*** (*t*), evaluated from the observed community state. These metrics are therefore instantaneous structural properties of the food web in a given month, not events directly recorded in the time series.

Because *ρ* and *t*_max_ are computed from the Jacobian at each monthly state, our analysis does not require observing the day-to-week phytoplankton response itself; we encoded that response in the local linearization. Nevertheless, monthly sampling implies that we average over within-month dynamics that might themselves include rapid bloom-crash episodes; this means that our metrics describe coarse-grained, monthly community configurations, and that fine-scale events between sampling dates are smoothed.

### Bioenergetic food-web model

Following [36], the dynamics of each guild are

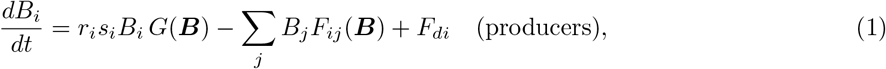

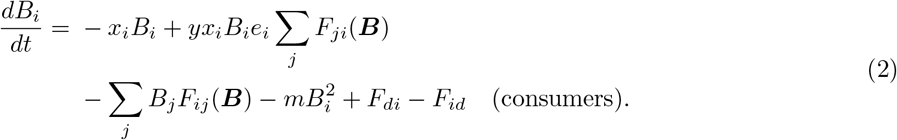

with

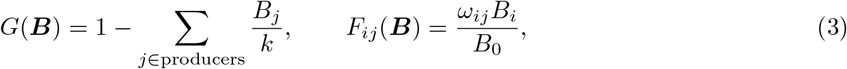

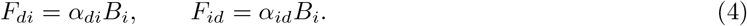

where *r*_*i*_ = the intrinsic growth rate of producer *i* (modulated by exudation fraction *s*_*i*_), *x*_*i*_ = the consumer metabolic rate, *y* = a maximum-consumption multiplier, *e*_*i*_ = assimilation efficiency, *ω*_*ij*_ = the (normalized) preference of *j* for resource *i, k* = the autotroph carrying capacity, *B*_0_ = the half-saturation constant, and *m* = a self-limitation coefficient. *F*_*di*_ and *F*_*id*_ are linear exchange terms with an auxiliary external compartment (node *d*) representing unresolved fluxes (advection, sedimentation, allochthonous inputs, benthic coupling). We took numerical parameter values from [36].

### Local linearization, mass-balance closure, and transient metrics

We did not assume that plankton communities sit at a long-term equilibrium. Instead, we treated each monthly observation as an operating point of a continuously forced system: at month *t* we evaluated the Jacobian ***J*** (*t*) at the observed state ***B***(*t*), so that interaction effects and stability properties vary with ecosystem state across successive observations. The auxiliary compartment *d* exists only to enforce instantaneous mass balance within the observed subnetwork: it absorbs net excess biomass and supplies biomass in deficit through the linear exchange terms above. We applied balance-enforcing closure, common in energetic food webs [38], solely to define the linearized operator ***J*** (*t*); it does not imply a literal biomass reservoir.

From ***J*** (*t*), we computed three complementary scalar descriptors of the response to a small perturbation ***x***(0): asymptotic stability, reactivity, and peak time.

### Asymptotic stability

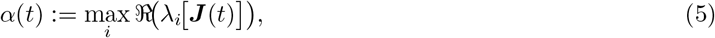

where *λ*_*i*_[***J*** (*t*)] denotes the *i*^*th*^ eigenvalue of ***J*** (*t*) and ℜ (·) extracts its real part. A fixed point is *linearly stable* if all eigenvalues have negative real parts, i.e., if *α*(*t*) < 0. In this case, small perturbations decay asymptotically and the rate of decay is governed by *α*(*t*); the quantity −1*/α*(*t*) defines the *asymptotic return time*, or relaxation time. After initial transients have decayed, the solution behaves as

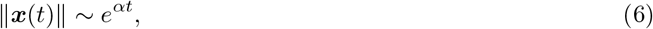

where *α* < 0 is the dominant (least stable) eigenvalue [20, 21], and ∥ ·∥denotes a vector norm (e.g., the Euclidean norm).

### Reactivity

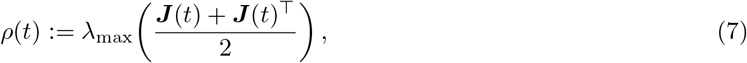

where *λ*_max_(·) denotes the largest eigenvalue of its argument and the superscript ⊤ denotes matrix transposition. For the symmetric part of ***J*** (*t*),

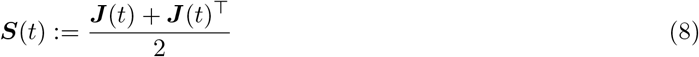

Because ***S***(*t*) is symmetric, all its eigenvalues are real and *λ*_max_(***S***(*t*)) is well defined. *ρ*(*t*) quantifies the system’s *reactivity* [26–28]: a positive *ρ* means that small perturbations can experience a transient growth in magnitude, even though the system is linearly stable overall (i.e., *α* < 0). Formally, *ρ*(*t*) is the maximum instantaneous growth rate of the squared norm of a perturbation ∥***x***(*t*) ∥^2^ [26], and therefore measures the extent to which the system can temporarily amplify small disturbances despite being asymptotically stable.

### Peak time

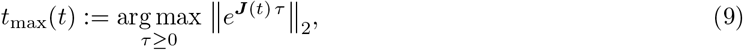

where ∥· ∥_2_ denotes the matrix 2-norm (spectral norm) induced by the Euclidean vector norm, and *e*^***J***(*t*) *τ*^ is the matrix exponential propagating the linearized dynamics over a time interval *τ*. This delay fixes when the worst-case overshoot *M*_max_ is realized. For a reactive (*ρ >* 0) and asymptotically stable (*α* < 0) equilibrium, the perturbation norm first grows up to *M*_max_ at *t*_max_ and then decays as exp[*α* (*t* −*t*_max_)]. As the measured abundances change over time, so do ***J*** (*t*) and all metrics computed from it. We give the full Jacobian decomposition (internal trophic terms *versus* closure-flux terms on the diagonal) in S1 Appendix, Section SI-A.

### Sensitivity analysis: temperature-dependent bioenergetic parameters

When using fixed bioenergetic parameters, warming should in reality modify those very parameters according to known thermal scaling [3, 4, 13]. As a sensitivity analysis (S1 Appendix, Section SI-G), we re-ran the entire pipeline with the producer growth rates *r*_*i*_ and consumer metabolic rates *x*_*i*_ scaled to the local monthly water temperature *T* (*t*) via a Boltzmann-Arrhenius factor,

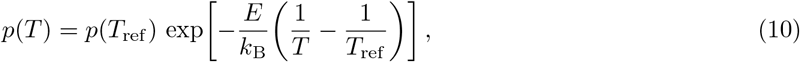

with reference temperature *T*_ref_ = 20 °C, activation energies *E*_*r*_ = 0.47 eV (autotroph growth), *E*_*x*_ = 0.65 eV (consumer metabolism), and *E*_*J*_ = 0.75 eV (ingestion, used only when *y* is also scaled), as is standard [4, 13]. We recomputed mass-balance closure terms *α*_*di*_, *α*_*id*_ consistently with the rescaled fluxes. We then compared the resulting reactivity time series (*ρ*_thermal_) to the original (*ρ*_fixed_) for each lake.

### Modelling time series with SARIMAX

To model the temporal dynamics of the response variable, we employed a two-step strategy that combined automated order selection with a seasonal autoregressive integrated moving average with exogenous regressors (SARIMAX) [37] framework. Our implementation included both systematic testing of stationarity and an iterative selection of exogenous predictors. The response variables were log *ρ*(*t*) and log *t*_max_(*t*); we treated asymptotic stability *α*(*t*) analogously but only report it in supplementary material because the patterns are weaker and lake-specific.

### Orthogonalization of environmental covariates in SARIMAX

Temperature and phosphate (PO_4_) might co-vary seasonally and over time (S2 Fig), which can induce collinearity when including multiple lags of both variables as exogenous regressors in SARIMAX. To improve coefficient stability and interpretability, we orthogonalized PO_4_ with respect to temperature using a residualization procedure.

For each lake, and after applying the same pre-processing used for the exogenous inputs, we built the lag block

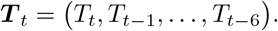

For each phosphate lag *l* ∈ {0, … , 6}, we then regressed *P*_*t−l*_ on ***T*** _*t*_ (including an intercept) and retained the residuals 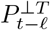. We fit the SARIMAX model using ***T*** _*t*_ and the residualized phosphate lags

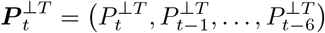

as exogenous covariates. In this formulation, temperature coefficients capture associations with temperature variability, while phosphate coefficients quantify associations with phosphate variability beyond that explained by temperature co-variation. As a sensitivity analysis, we repeated the procedure in S1 Appendix, Section SI-F by swapping the roles of the covariates (temperature residualized on the PO_4_ lag block).

### Preprocessing and stationarity testing

We assessed stationarity of each series using the augmented Dickey–Fuller test [39]. The covariates we considered included temperature and phosphate concentration (PO_4_). We differenced non-stationary series until the null hypothesis of a unit root could be rejected when the type I error estimate exceeded 0.05. After determining if it was necessary (S1 Appendix, Section SI-D), we applied a logarithmic transformation to the response variable, making sign corrections when needed, followed by differencing if required. Additionally, we tested seasonal unit roots using 12-month differences, and set the corresponding seasonal integration order *D* to either 0 or 1 as appropriate.

### Construction of exogenous variables

We built exogenous regressors from both contemporaneous and lagged values of temperature and PO_4_, with lags up to a maximum of 6 months. This allowed the model to capture delayed effects of environmental covariates on the response. We then subjected all candidate exogenous variables to a multicollinearity reduction step based on the variance inflation factor [40]. We only retained predictors with variance inflation < 5.0.

#### Step 1: model order selection

We first did an automatic autoregressive integrated moving-average (ARIMA) order selection procedure, that evaluated a set of candidate (*p, d, q*) × (*P, D, Q*)_*m*_ specifications, where *p, d, q* = the non-seasonal autoregressive, differencing, and moving-average orders, *P, D, Q* = their seasonal counterparts, and *m* the length of the seasonal cycle. We chose the optimal orders by minimizing Akaike’s information criterion (AIC) [41].

#### Step 2: estimation via SARIMAX

Given the selected orders, we re-estimated the model using the SARIMAX specification. This allows the inclusion of exogenous regressors and a richer set of diagnostic tools (S1 Appendix, Section SI-D). The first step ensures that the lag orders are chosen in a data-driven and statistically principled way, avoiding *ad hoc* specification. The second step leveraged the more general SARIMAX estimation framework to incorporate exogenous information and calculate formal model diagnostics.

#### Model evaluation

We compared competing specifications using Akaike’s information criterion. We retained the model with the lowest AIC among all tested autoregressive orders, lag structures, and exogenous subsets as the top-ranked specification. This procedure balanced predictive accuracy with parameter minimisation, while accounting for potential multicollinearity and seasonal structure. Because *ρ*(*t*) and *t*_max_(*t*) are model-derived rather than directly observed, the resulting SARIMAX coefficients quantify how environmental drivers covary with the model-implied transient metrics under the Boit et al. [36] parameterization; we used the Boltzmann-Arrhenius sensitivity analysis (above) to assess robustness to this parameterization.

## Results

### Lake food webs occupy an asymptotically stable yet reactive regime modified by warming

Across all lakes, water temperature showed a long-term increase over the study period, while phosphate concentrations declined substantially in some systems and modestly or not at all in others (Fig. 3; Table S2). Despite this heterogeneity in nutrient trajectories, every lake-month sat in a regime that was simultaneously stable (*α* < 0) and reactive (*ρ >* 0) (Fig S1). Over time, lakes tended to drift toward less reactive but also less stable configurations, a co-evolution of *ρ* and *α* consistent with theoretical expectations for systems near regions of structural change [29].

### Warming coherently raises reactivity and lengthens the time to peak response

A direct visual assessment (Fig S5) shows that in each lake, monthly temperature is generally positively associated with log *ρ* and, more weakly and inconsistently, with log *t*_max_; phosphate associations are weaker and lake-specific. To quantify these associations rigorously, we estimated SARIMAX models with temperature lags and phosphate residualized on temperature 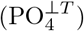; for each lake, we report the average coefficient across the temperature lags receiving the strongest statistical support (Fig. 4; full lag-wise estimates in Fig S9).

**Fig 4.**
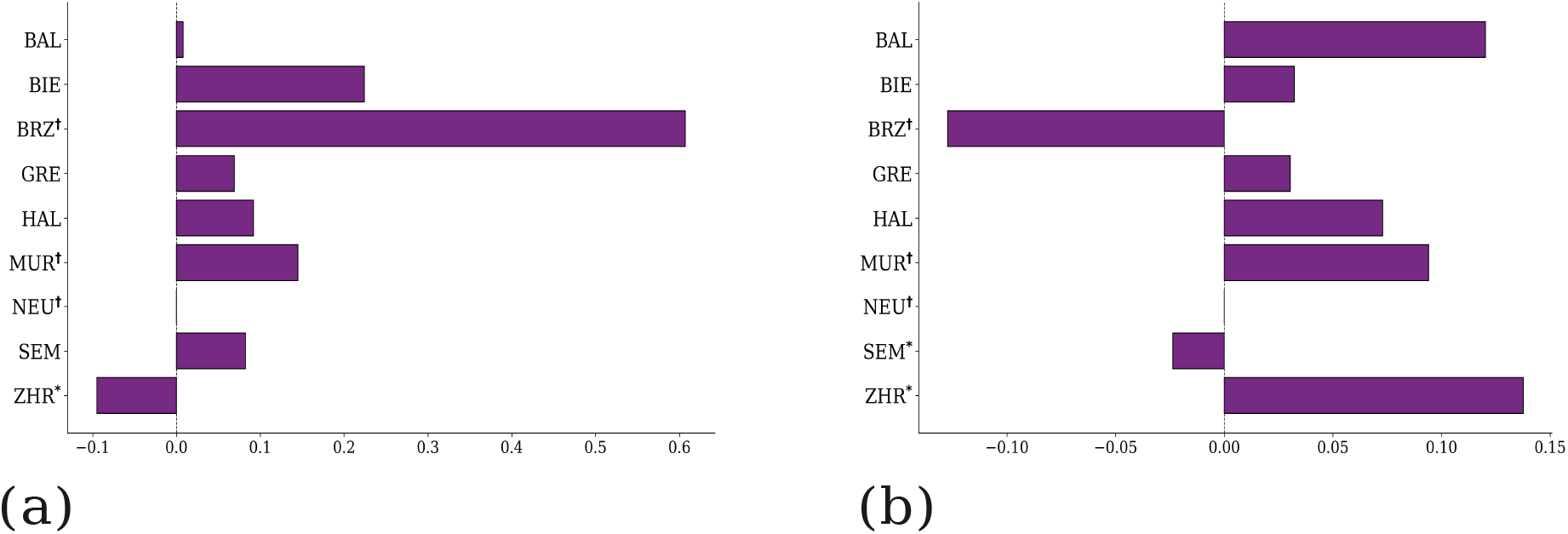
Temperature effects on transient metrics from SARIMAX models with orthogonalized phosphate. Lake-specific coefficients for (a) log(*ρ*) and (b) log(*t*_max_), estimated from SARIMAX models that include temperature lags and phosphate residualized with respect to the temperature lag block 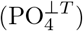. For each lake, bars show the average temperature coefficient across lags with the strongest statistical support (full lag-specific estimates and uncertainty: S9 Fig). Lake acronyms: BAL (Baldegg), BIE (Biel), BRZ (Brienz), GRE (Greifen), HAL (Hallwil), MUR (Murten), NEU (Neuenburg), SEM (Sempach), ZHR (Zurich). Asterisks (*) flag lakes where residual diagnostics indicated potential misspecification; daggers (†) flag lakes where the absolute PO_4_ change is < 5% of the maximum observed variation. Coefficients refer to the temperature predictor.

Two patterns emerge across lakes. First, temperature coefficients on log *ρ* are predominantly positive (Fig. 4a): warmer conditions are associated with stronger short-term amplification, in the direction predicted by thermal scaling of consumer-resource interaction strengths. Translating these coefficients into magnitudes, a 1 °C rise in mean temperature corresponds to a ∼ 10% higher reactivity on average across lakes. Second, temperature coefficients on log *t*_max_ are also mostly positive (Fig. 4b), meaning that warming is associated with delayed peak amplification (indicating slower but more prolonged transient excursions from equilibrium). A few lake-specific exceptions occur and we interpret these with caution where residual diagnostics flag potential model misspecification (asterisks).

In contrast, residualized phosphate effects 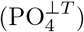 on either metric are rarely distinguishable from zero and lack a consistent sign across lakes (Fig S6). Once we accounted for temperature-phosphate collinearity, nutrient variation alone is not an equally informative predictor of transient amplification or its timing. These conclusions are robust to alternative collinearity treatments, residualizing temperature on phosphate, or using raw PO_4_ (Fig S4), and to using a fully temperature-scaled bioenergetic parameterization (Section ; Fig S12).

### Asymptotic stability is lake-specific and has weaker associations

The same SARIMAX framework applied to log(−*α*) yields coefficients whose sign and magnitude vary across lakes, with no coherent cross-lake pattern (Fig S7). Because our focus is transient dynamics, and because *α* is also more vulnerable to assumptions about the closure compartment (see Section), we report these results in the supplementary material only.

### Consumer biomass tracks reactivity; predator biomass tracks *t*_max_

Because we fixed the trophic topology across time, month-to-month changes in *ρ* and *t*_max_ necessarily reflect changes in the biomass distribution across guilds. We therefore correlated Δ*ρ* and Δ*t*_max_ with Δ biomass for each guild within each lake (Fig. 5).

**Fig 5.**
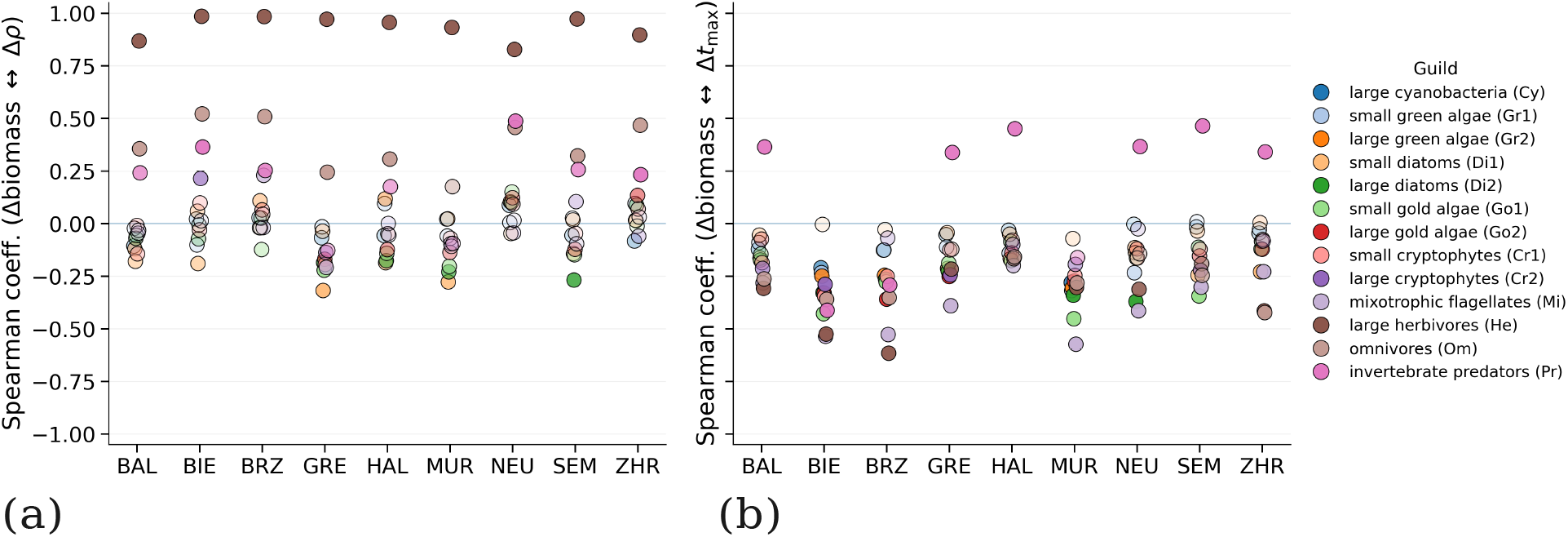
Guild-level correlates of transient response magnitude and timing across lakes. Each point indicates the Spearman correlation between month-to-month changes in guild biomass (Δ biomass) and changes in transient-response metrics computed from the Jacobian time series. **(a)** Correlations between Δ biomass and changes in the time to peak response (Δ*t*_max_). **(b)** Correlations between Δ biomass and changes in reactivity (Δ*ρ*). Colours identify guilds; the horizontal line indicates zero correlation. Marker transparency reflects the strength of statistical evidence for a correlation, with more opaque symbols corresponding to higher evidence. Lake acronyms: BAL (Lake Baldegg), BIE (Lake Biel), BRZ (Lake Brienz), GRE (Lake Greifen), HAL (Lake Hallwil), MUR (Lake Murten), NEU (Lake Neuenburg), SEM (Lake Sempach), and ZHR (Lake Zurich).

Two patterns emerged consistently across lakes. Δ*ρ* was most strongly and most consistently correlated with Δ biomass of large herbivores, with secondary contributions from omnivores and large invertebrate predators (Fig. 5b); primary-producer guilds had much weaker, lake-specific associations (Fig. 5b). Δ*t*_max_ was predominantly negatively correlated with Δ biomass for most guilds (i.e., increases in biomass tend to coincide with earlier peaks) (Fig. 5a), with one systematic exception: invertebrate predators had positive correlations across all lakes, indicating that increases in predator biomass pushed the peak amplification later. This pattern is consistent with the theoretical expectation that *ρ* is dominated by a few strongly coupled, structurally central interaction channels [27, 34], and with the ecological fact that large zooplankton grazers and intraguild predators occupy precisely those channels in plankton food webs.

### Robustness to temperature-dependent bioenergetic parameters

We estimated the above associations with bioenergetic parameters fixed to the values from [36], but warming is expected to modify those parameters themselves through thermal scaling [3, 4, 13]. To test whether our central finding survives this internal feedback, we re-ran the entire pipeline with *r*_*i*_ and *x*_*i*_ rescaled to monthly water temperature via the Boltzmann-Arrhenius relation (Eq. 10; Methods). Across the 2,949 lake-months for which we could compute both versions, the rank ordering of reactivity was essentially preserved (Spearman *ρ >* 0.94 in every lake, *ρ* = 0.988 globally), and the cross-method correlation was high also in absolute value (Pearson *r >* 0.88 per lake, *r* = 0.97 globally; S1 Appendix, Section SI-G, S5 Table). Absolute values of *ρ* were systematically lower under thermal scaling because most lake temperatures were below the reference *T*_ref_ = 20 °C, which down-regulates rates; however, the sign and lake patterns of warming-driven reactivity change was preserved. The mechanistic temperature-reactivity association is therefore not an artifact of fixing bioenergetic parameters.

## Discussion

We provide model-assisted empirical evidence that long-term warming has systematically reshaped the transient response of nine lake plankton food webs to small perturbations. After controlling for autocorrelation, seasonality, lagged effects, and temperature-phosphate collinearity, higher water temperature was consistently associated with both higher reactivity and longer time to peak amplification across lakes. In contrast, phosphate effects on these transient metrics were weak and lake-specific once we accounted for temperature variation. Together, these patterns indicate that warming is associated with stronger and more prolonged transient excursions from the local equilibrium, even though every observed community state remained asymptotically stable.

### Connecting transient metrics to thermal ecology

Our central observation that warming raises reactivity and lengthens the time to peak response fits expectations from theoretical thermal ecology [3, 4, 13, 15, 33]. Temperature does not act on a single rate; it modifies metabolic demand, autotroph production, attack rates and handling times with rate- and taxon-specific sensitivities [3, 14, 19]. These changes can alter per-capita consumer-resource interaction strengths, and stronger off-diagonal interaction terms are expected to increase the dominant eigenvalue of the symmetric part of the Jacobian, i.e., reactivity [26, 27, 34]. Our SARIMAX results provide a multidecadal field-scale signature consistent with this mechanism, complementing microcosm and modelling evidence [15, 33, 42]. The robustness of the warming-reactivity association under explicit Boltzmann-Arrhenius scaling of the bioenergetic parameters (S1 Appendix, Section SI-G) further indicates that this signature is not an artifact of fixing interaction strengths at reference temperatures.

The longer time-to-peak response under warming is more subtle. Warming compresses several individual-level timescales, but transient amplification is shaped by the geometry of the linear operator ***J*** rather than by the speed of any single process. When interaction strengths and non-normality change, a small perturbation can reach a larger peak after a longer delay, even if all perturbations eventually decay [26, 27]. This is consistent with the positive association between temperature and *t*_max_ we observed.

### Why consumer guilds, not autotrophs, mediate the change

Reactivity is expected to be sensitive to strong interaction terms in the Jacobian [27, 34]. In planktonic food webs, such terms are likely to occur along consumer-resource and intraguild predation links involving large herbivores, omnivores, and predators. The strong consumer-driven covariation between Δ biomass and Δ*ρ* (Fig. 5b), together with the predator-specific positive covariation with Δ*t*_max_ (Fig. 5a), is mechanistically consistent with this interpretation. Warming can shift these consumer-mediated pathways by altering generation times, top-down pressure, and asymmetric rate scaling between consumers and producers [14, 15, 33]. This varies interaction strengths in the same consumer and predator channels that our analysis flags.

### Implications for early-warning indicators and management

Critical slowing down has been proposed as an early-warning indicator of impending dynamical regime shifts [43], and elevated reactivity has been similarly suggested as a precursor to loss of stability [27]. We do not claim that any of the lakes we examined are approaching a tipping point, nor that we can identify a threshold. But our results do show that the transient dynamical regime has shifted coherently across lakes: communities now amplify perturbations more strongly and remain in displaced configurations for longer than they did decades ago. This raises ecological sensitivity to any subsequent disturbance – heat waves, stratification anomalies, sudden nutrient pulses, or biological invasions – during the window in which the system is still relaxing.

The contrast with phosphate is informative for management. Re-oligotrophication policies have substantially reduced nutrient loads in most of the studied lakes, and our analysis confirms that this nutrient reduction has not, on its own, generated a coherent reactivity signal. In contrast, warming has.

Nutrient-focused management is necessary, but apparently not sufficient to compensate for warming-driven changes in transient dynamics; explicit monitoring of and intervention on warming-related drivers of transient risk (heat-wave occurrence, stratification regimes, disturbance frequency) is therefore warranted in parallel.

### Limitations and outlook

#### Model-derived metrics carry uncertainty not propagated into SARIMAX

Our two-layer design treats *ρ*(*t*) and *t*_max_(*t*) as response variables in the SARIMAX layer, even though they are derived from a bioenergetic model whose parameters [36] are themselves uncertain. We do not formally propagate this parameter uncertainty into our SARIMAX confidence intervals, and our reported coefficients should be interpreted as describing how warming relates to model-implied transient metrics, not as estimates of how warming relates to a directly observed quantity. Three considerations partly mitigate this concern: (*i*) the cross-lake coherence of the warming-reactivity association argues against a parameterization-specific artifact; (*ii*) the Boltzmann-Arrhenius sensitivity analysis (Section and S1 Appendix, Section SI-G) preserves the qualitative result when the bioenergetic parameters themselves are temperature-dependent; and (*iii*) the orthogonalization strategy isolates temperature signals from phosphate signals.

#### Monthly resolution and short-term phytoplankton dynamics

Phytoplankton can respond to perturbations on day-to-week timescales [6, 30]. Our metrics are not direct measurements of those responses; they are instead properties of the local linearization at each monthly state. Monthly sampling therefore averages over within-month fluctuations and might smooth out genuinely fast events. This temporal coarse-graining most likely explains the weak, lake-specific correlations between primary-producer biomass and transient metrics (Fig. 5): rapid producer turnover between sampling dates is invisible to our observation grid, while slower-changing consumer guilds aggregate more cleanly.

Higher-frequency sampling would test this directly.

#### Asymptotic stability and the closure compartment

The asymptotic stability *α* depends on diagonal Jacobian entries that include both internal trophic terms (self-limitation, intraspecific competition) and contributions from the external closure flux. Closure terms could plausibly change systematically with warming and oligotrophication, so part of any apparent *α* trend might reflect closure dynamics rather than internal community self-regulation. Because we have de-emphasized *α* throughout (focusing on *ρ* and *t*_max_, which are governed primarily by the symmetric off-diagonal structure), this concern does not threaten our central conclusion. A formal decomposition of the diagonal into internal and closure contributions, and a sensitivity analysis on the closure parameterization, are useful next steps.

#### Fixed topology

We fixed the trophic topology across time; this is a deliberately conservative choice that lets us isolate state-dependence from network change but obviously underestimates real interaction-network plasticity over decades.

#### Causality and unmeasured drivers

Our analysis is correlative with respect to environmental forcing. We considered both temperature and phosphate, but additional drivers (mixing regime, light, nitrogen, species composition, invasions) could plausibly shape the dynamics we report. The cross-lake coherence of the warming signal, the robustness checks against collinearity, the consistent guild-level mechanism, and the resilience of the central result under Boltzmann-Arrhenius scaling together support, but do not prove, a temperature-driven mechanism.

## Acknowledgments

GVC acknowledges the JRC Exploratory Research Programme, which supported the Exploratory Research Project “REloading the Anthropocene to Decode Mass Extinction (README)”. We are grateful to Francesco Pomati and his research group at Eawag for their constructive comments and valuable discussions.

## Supporting information

**S1 Appendix. Supplementary methods, tables, and figures**.

### SI-A. Lakes, guilds, and percent changes

#### Nodes and lakes

**S1 Table.**
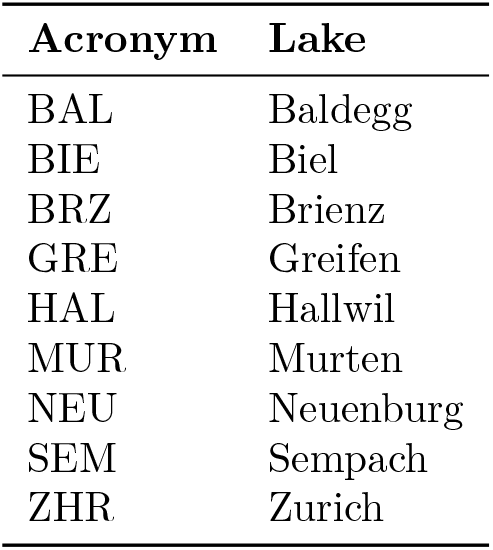
List of Swiss lakes included in the study with corresponding acronyms used throughout the text and figures.

**S2 Table.**
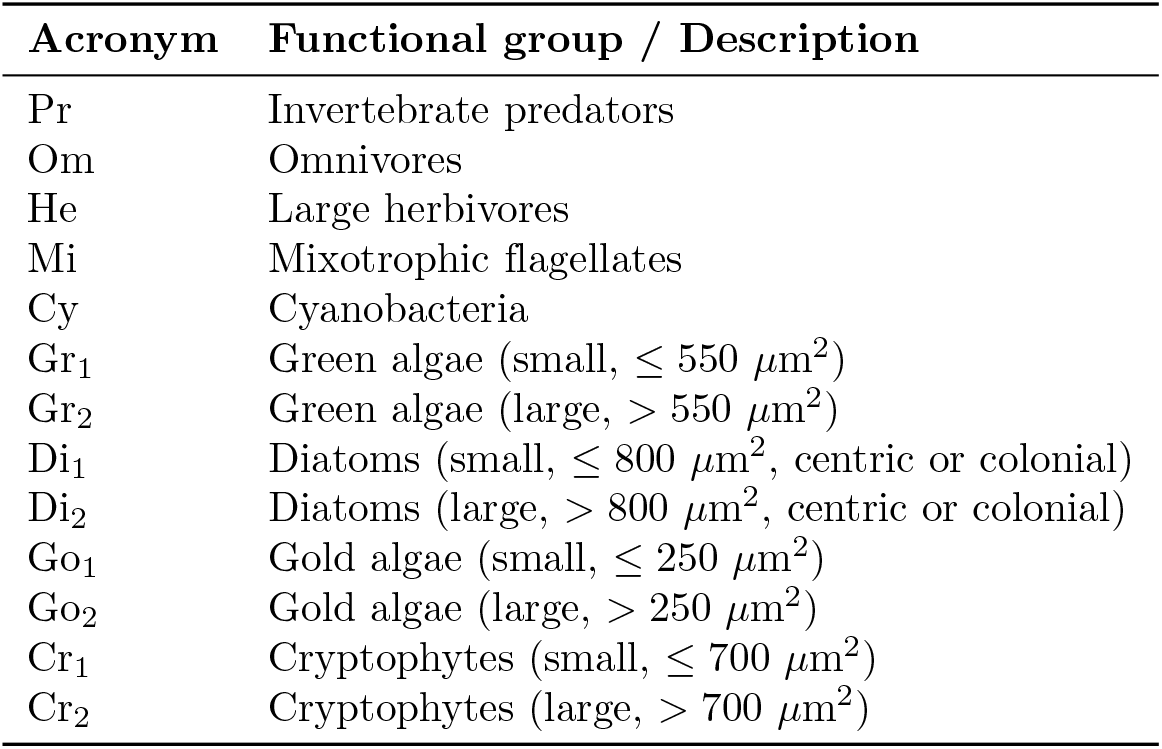
Functional groups included in the planktonic food-web model with their acronyms and definitions.

| Acronym | Functional group / Description |
| --- | --- |
| Pr | Invertebrate predators |
| Om | Omnivores |
| He | Large herbivores |
| Mi | Mixotrophic flagellates |
| Cy | Cyanobacteria |
| Gr<sub>1</sub> | Green algae (small, ≤ 550 μm<sup>2</sup>) |
| Gr<sub>2</sub> | Green algae (large, > 550 μm<sup>2</sup>) |
| Di<sub>1</sub> | Diatoms (small, ≤ 800 μm<sup>2</sup>, centric or colonial) |
| Di<sub>2</sub> | Diatoms (large, > 800 μm<sup>2</sup>, centric or colonial) |
| Go<sub>1</sub> | Gold algae (small, ≤ 250 μm<sup>2</sup>) |
| Go<sub>2</sub> | Gold algae (large, > 250 μm<sup>2</sup>) |
| Cr<sub>1</sub> | Cryptophytes (small, ≤ 700 μm<sup>2</sup>) |
| Cr<sub>2</sub> | Cryptophytes (large, > 700 μm<sup>2</sup>) |

#### Summary of percent changes in ecological and environmental metrics across Swiss lakes

Table S3 summarizes the percent change between the first and the last 36-month windows of observation for the main ecological and environmental indicators in each lake.

**S3 Table.** Changes (Δ and %) for reactivity (*ρ*), asymptotic stability (*α*), PO_4_ (*µ*g/L), and temperature (°C). Values are differences between the averages of the first 36 and last 36 valid observations for each variable.

| Lake | $\rho$ $\Delta$ | $\rho$ % | $\alpha$ $\Delta$ | $\alpha$ % | $\text{PO}_4$ $\Delta$ [$\mu\text{g/L}$] | $\text{PO}_4$ % | Temp. $\Delta$ [$^\circ\text{C}$] | Temp. % |
| --- | --- | --- | --- | --- | --- | --- | --- | --- |
| HAL | -50.74 | -18.93 | 0.0095 | 30.47 | -0.14 | -99.03 | 1.06 | 13.12 |
| NEU | -22.34 | -8.02 | 1.30 | 67.75 | -0.00047 | -10.07 | 0.18 | 2.68 |
| ZHR | -153.24 | -43.68 | 0.0041 | 17.88 | -0.051 | -70.89 | 1.38 | 18.91 |
| BRZ | -15.63 | -20.10 | 1.67 | 65.76 | -0.0040 | -79.23 | 0.36 | 6.22 |
| BAL | -54.85 | -16.85 | 0.015 | 30.80 | -0.14 | -92.65 | 1.25 | 19.63 |
| GRE | -2.00 | -0.79 | 0.015 | 26.20 | -0.078 | -58.11 | 0.38 | 4.61 |
| MUR | 89.88 | 69.71 | 1.11 | 25.01 | -0.0023 | -50.01 | 1.32 | 17.39 |
| SEM | -44.41 | -10.88 | 0.083 | 62.63 | -0.084 | -86.44 | 0.71 | 10.21 |
| BIE | -0.13 | -0.065 | 1.26 | 32.14 | -0.017 | -73.47 | 1.05 | 14.05 |

### SI-B. Jacobian matrix

The Jacobian matrix 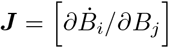 describes the local dynamics of biomass change with respect to species interactions:

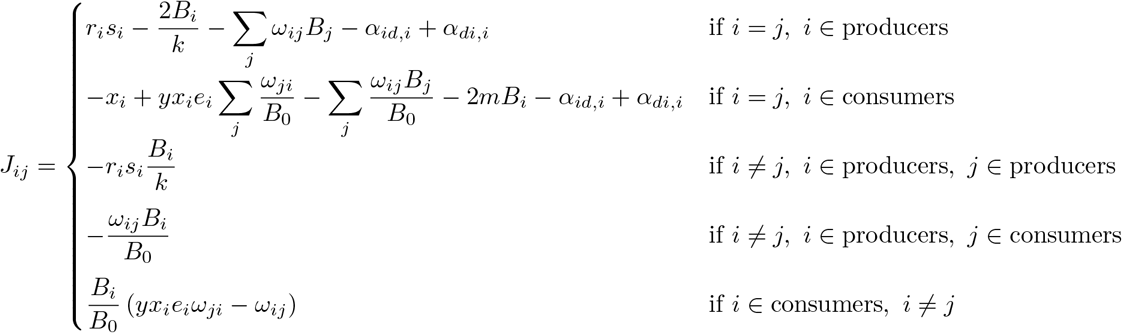

with *ω*_*ij*_ = *a*_*ij*_*/K*_in_(*j*) the normalized interaction strength. Parameters include metabolic loss *x*_*i*_, assimilation efficiency *e*_*i*_, self-limitation *m*, closure flows *α*_*di,i*_ and *α*_*id,i*_, and saturation constant *B*_0_. The constant *y* is an efficiency multiplier. Producers have *s*_*i*_ *>* 0 and no incoming links (*K*_in_ = 0).

The diagonal entries thus combine internal contributions (self-limitation −2*B*_*i*_*/k* for producers, −2*mB*_*i*_ for consumers; metabolic loss −*x*_*i*_; trophic feedback through *ω*_*ij*_*B*_*j*_) and closure contributions (−*α*_*id,i*_ + *α*_*di,i*_).

### SI-C. Correlation between stability metrics

**S4 Table.**
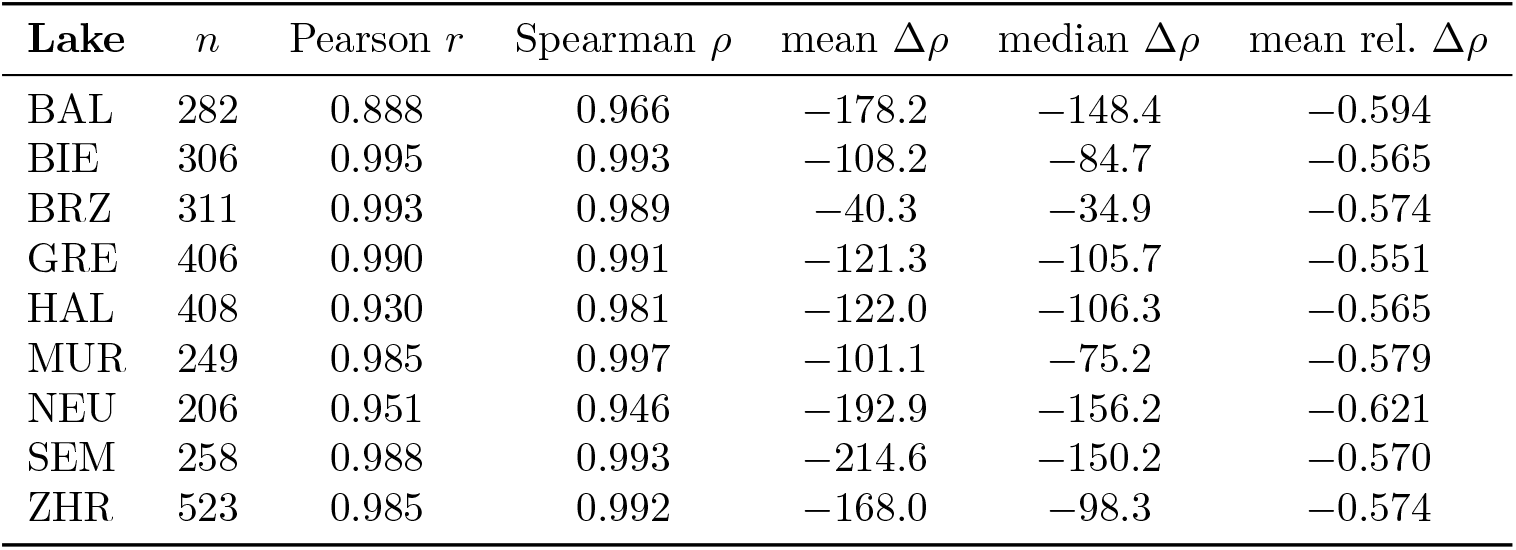
Mean Pearson correlations across all lakes between reactivity (*ρ*), asymptotic stability (*α*), and time to peak response (*t*_max_).

### SI-D. SARIMAX model details

Before estimating a SARIMAX (seasonal autoregressive integrated moving average with exogenous regressors) model, it is important to evaluate whether a logarithmic transformation of the dependent variable is appropriate [37, 44]. The log-transformation can reduce skewness, stabilize variance, and improve the linearity and homoscedasticity assumptions required by regression- and ARIMA-type models. To assess this, we performed a set of diagnostic checks on the candidate variable:

- **Distributional properties**. Skewness and kurtosis were computed and histograms were compared before and after log-transformation. A log-transform is expected to reduce asymmetry and excess kurtosis, bringing the distribution closer to normality.
- **Variance stability**. The rolling standard deviation (with a fixed window size) was calculated for the original and the log-transformed series. If the rolling standard deviation of the logged series is more stable, the log transformation helps in controlling heteroskedasticity over time.
- **Homoscedasticity tests**. The Breusch-Pagan test was applied to both the original and log-transformed data. A higher *p*-value after transformation suggests reduced heteroskedasticity.

We used a logarithmic transformation when at least two of the above diagnostics showed improvement (e.g., reduced skewness, reduced kurtosis, stabilized variance, or lower heteroskedasticity). This diagnostic step is important because SARIMAX, like other regression-ARIMA models, assumes approximately homoscedastic and well-behaved residuals. By ensuring that the dependent variable has more stable variance and a distribution closer to normality, we improved the reliability of parameter estimates and the interpretability of the model.

#### Residual diagnostics

To assess the adequacy of the fitted SARIMAX models, we computed several residual diagnostics on the standardized residual series. A well-specified time-series model should produce residuals that approximate white noise (zero mean, constant variance, no remaining autocorrelation):

- **Mean and variance**. We checked that the residual mean was close to zero and compared the standard deviation across lakes.
- **Durbin-Watson statistic**. Used to detect first-order autocorrelation in residuals; values close to 2 indicate absence of serial correlation [45].
- **Ljung-Box test**. Applied over lags 1, … , *L*(= 12), with reported minimum, mean, and median type-I error estimates. Large *p*-values suggest no residual correlation [46].
- **Anderson-Darling test**. Used to assess the null hypothesis of normality of residuals [47].
- **ARCH test**. Tests for autoregressive conditional heteroskedasticity in residuals; evidence for this property suggests volatility clustering not captured by the SARIMAX specification [48].

Rather than reporting full diagnostic tables, we summarized residual behavior using a conservative *soft-flag* scheme intended solely as a visual aid. Lakes marked with an asterisk in figures correspond to cases where residual diagnostics consistently indicated departures from standard modeling assumptions across multiple tests and lags. Specifically, a lake was flagged when at least one of the following patterns was observed: (i) **residual autocorrelation:** Ljung-Box diagnostics showed systematically low *p*-values across tested lags; (ii) **conditional heteroskedasticity:** ARCH tests exhibited persistently low average *p*-values; or (iii) **non-normality with shape deviation:** the Anderson-Darling test indicated non-Gaussian residual structure with clear departures in skewness or kurtosis. These flags are not used for model selection or inference; they only highlight cases where residual assumptions warrant additional caution.

### SI-E. Lake-level distributions and bivariate plots

**S1 Fig.**
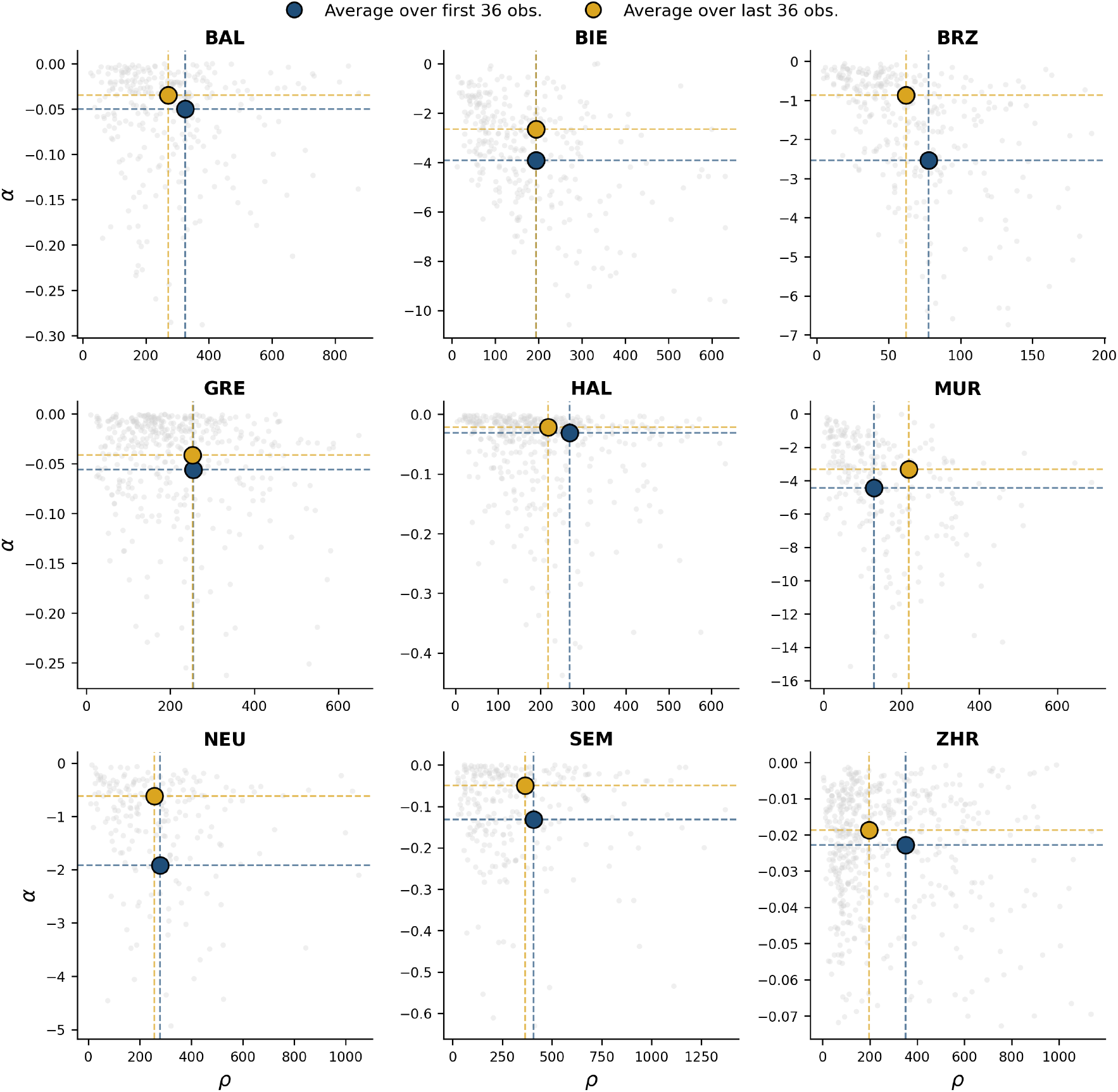
Bivariate distributions of reactivity (*ρ*) and asymptotic stability (*α*) for each lake. Points represent individual monthly estimates, while dark blue and yellow markers denote the centroids (mean values) of the first and last 36 valid observations, respectively. Dashed lines indicate the corresponding average values along each axis.Lake acronyms denote: BAL (Lake Baldegg), BIE (Lake Biel), BRZ (Lake Brienz), GRE (Lake Greifen), HAL (Lake Hallwil), MUR (Lake Murten), NEU (Lake Neuenburg), SEM (Lake Sempach), and ZHR (Lake Zurich).

**S2 Fig.**
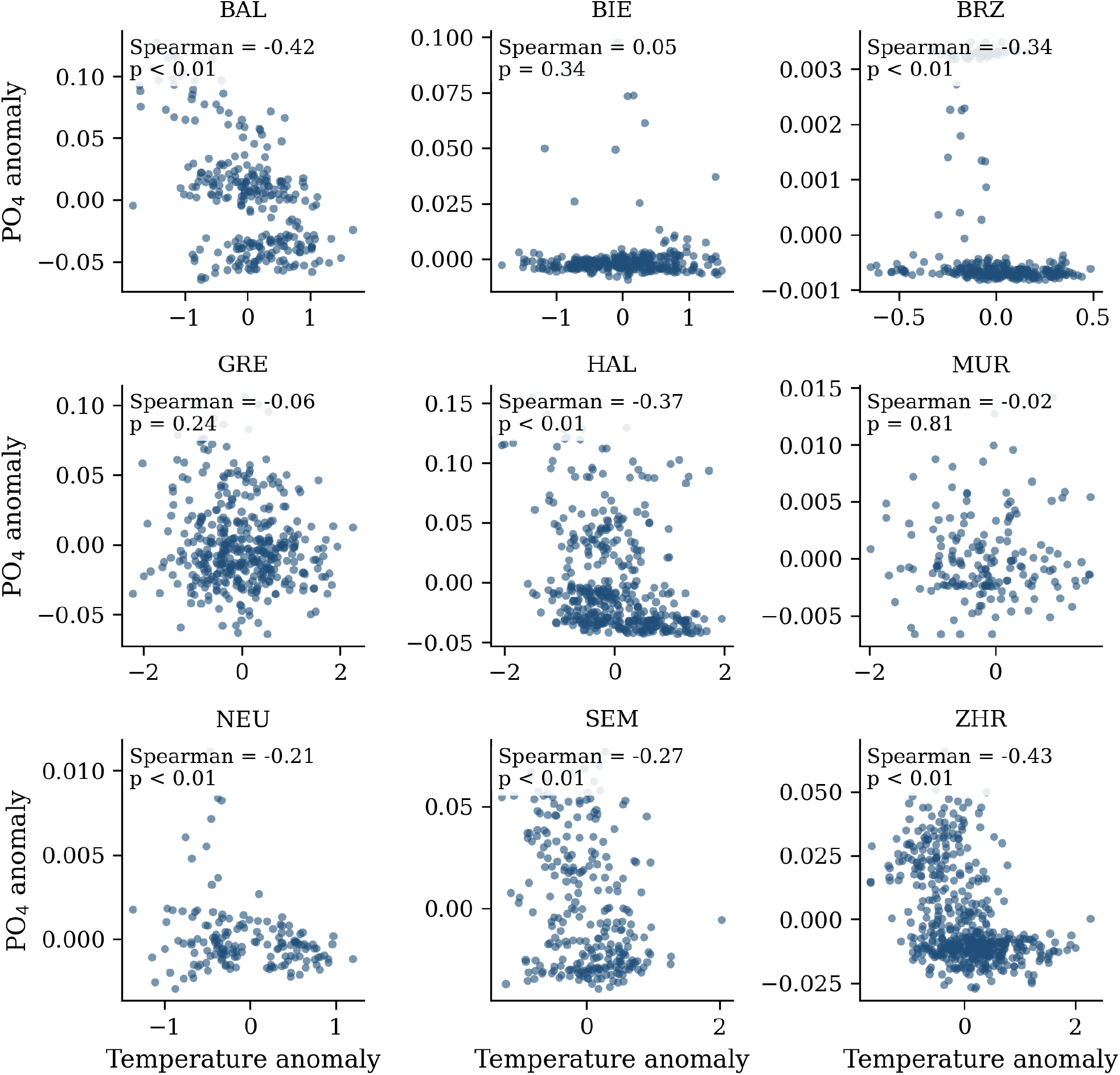
Scatterplots of temperature versus PO_4_ monthly anomalies for nine lakes. We calculated anomalies by removing the long-term monthly mean. Panels report Spearman’s rank correlation coefficient and type I error estimate. Lake acronyms denote: BAL (Lake Baldegg), BIE (Lake Biel), BRZ (Lake Brienz), GRE (Lake Greifen), HAL (Lake Hallwil), MUR (Lake Murten), NEU (Lake Neuenburg), SEM (Lake Sempach), and ZHR (Lake Zurich).

**S3 Fig.**
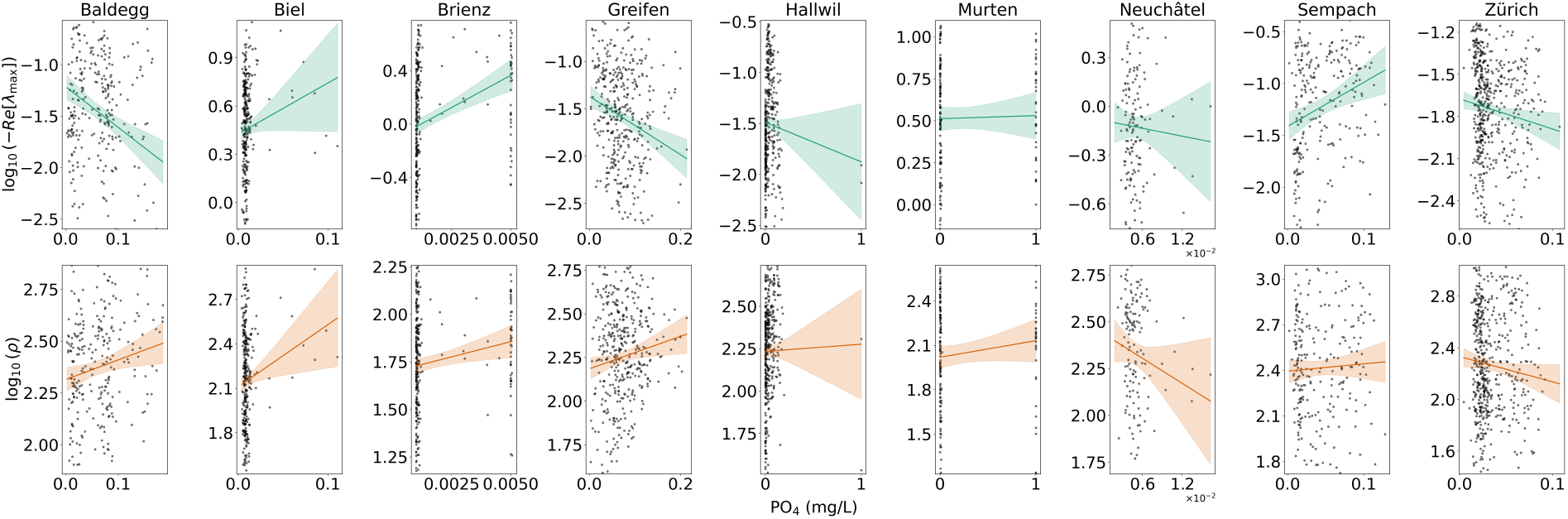
For each lake, relationship between phosphate concentration (PO_4_, µg/L) and two stability metrics: top row log_10_(−Re[*λ*_max_]) (teal) and bottom row log_10_(*ρ*) (orange). Points show monthly values; solid lines represent ordinary least squares fits with 95% confidence intervals.

**S4 Fig.**
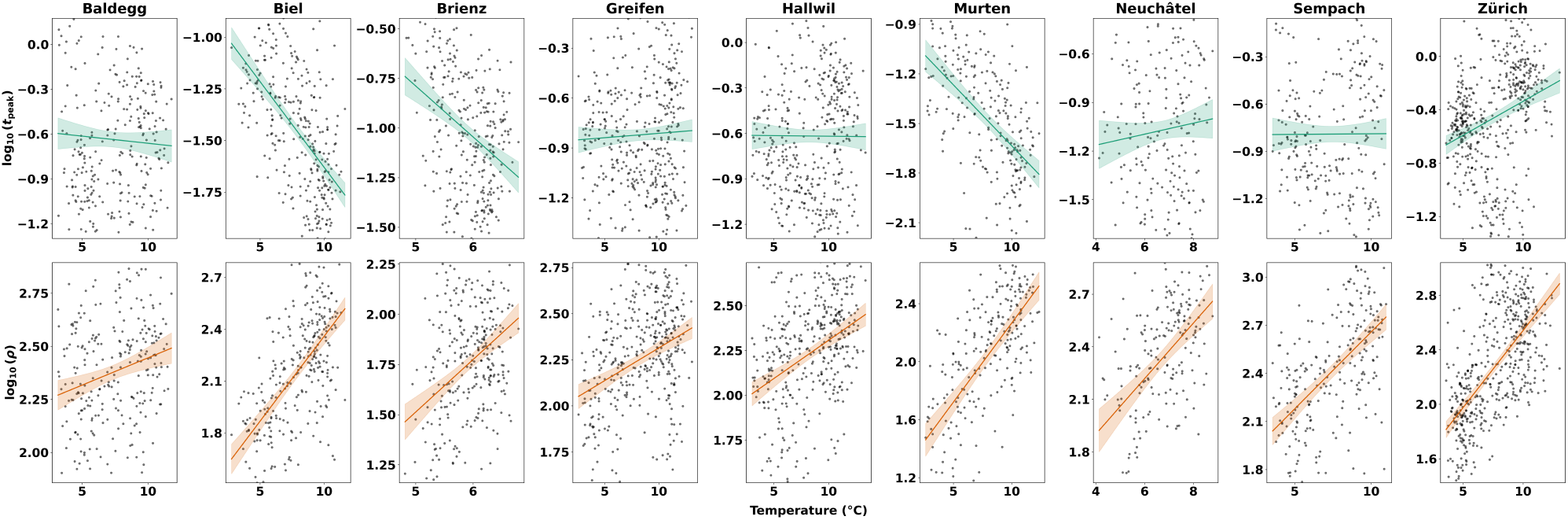
For each lake, monthly water temperature (°C) is related to two transient-response metrics: *t*_max_ (top row; log_10_(*t*_max_), teal) and *ρ* (bottom row; log_10_(*ρ*), orange). Points denote monthly observations; solid lines are ordinary least-squares fits and shaded bands indicate 95% confidence intervals.

### SI-F. SARIMAX results: alternative orthogonalization and PO_4_ effects

**S5 Fig.**
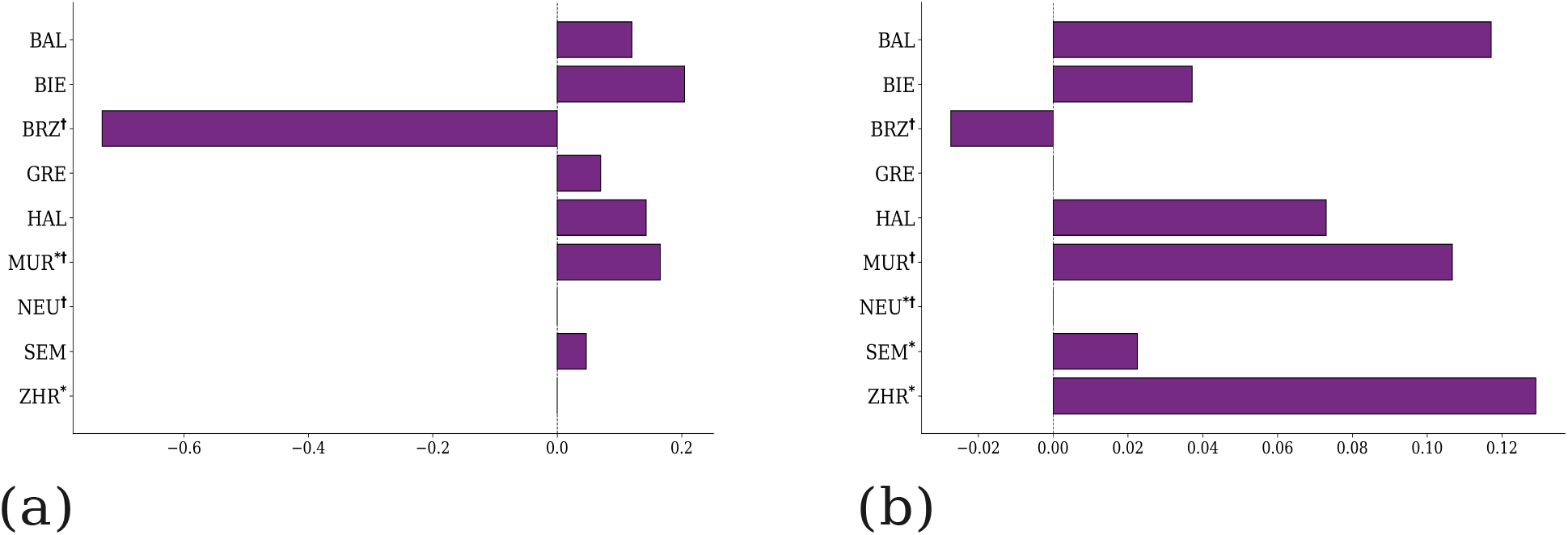
Temperature effects from SARIMAX models with temperature residualized on phosphate. Lake-specific coefficients for (a) log(*ρ*) and (b) log(*t*_max_) from SARIMAX models including phosphate lags and temperature residualized with respect to the phosphate lag block 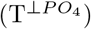. Bars: average coefficient across lags with strongest statistical support. Lake acronyms denote: BAL (Lake Baldegg), BIE (Lake Biel), BRZ (Lake Brienz), GRE (Lake Greifen), HAL (Lake Hallwil), MUR (Lake Murten), NEU (Lake Neuenburg), SEM (Lake Sempach), and ZHR (Lake Zurich). An asterisk (*) denotes lakes where residual diagnostics failed, and a dagger (†) denotes lakes where the absolute PO_4_ change is < 5% of the maximum observed variation.

**S6 Fig.**
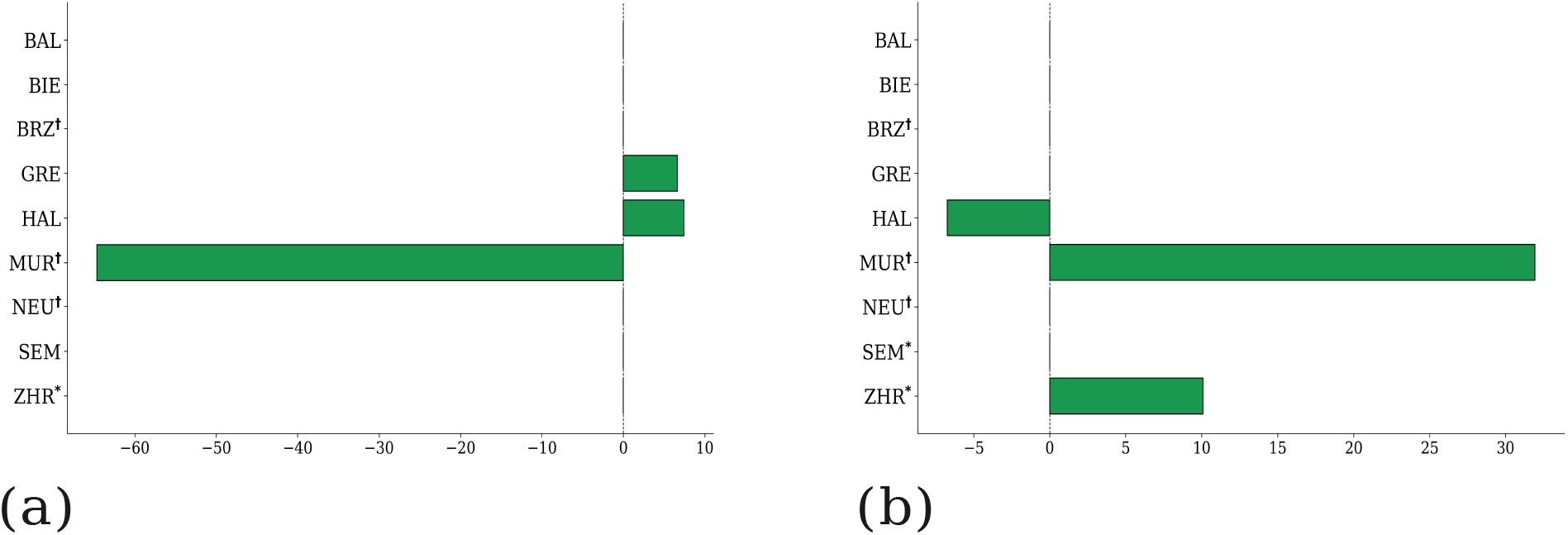
PO_4_ effects from SARIMAX models with orthogonalized phosphate. Lake-specific coefficients for the phosphate predictor for (a) log(*ρ*) and (b) log(*t*_max_), estimated from SARIMAX models including temperature lags and phosphate residualized with respect to the temperature lag block For each lake, bars summarize the average phosphate coefficient across lags that show the strongest statistical support in that lake. Lake acronyms denote: BAL (Lake Baldegg), BIE (Lake Biel), BRZ (Lake Brienz), GRE (Lake Greifen), HAL (Lake Hallwil), MUR (Lake Murten), NEU (Lake Neuenburg), SEM (Lake Sempach), and ZHR (Lake Zurich). An asterisk (*) denotes lakes where residual diagnostics failed, and a dagger (†) denotes lakes where the absolute PO_4_ change is < 5% of the maximum observed variation. All reported coefficients refer to the 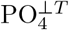 predictor.

**S7 Fig.**
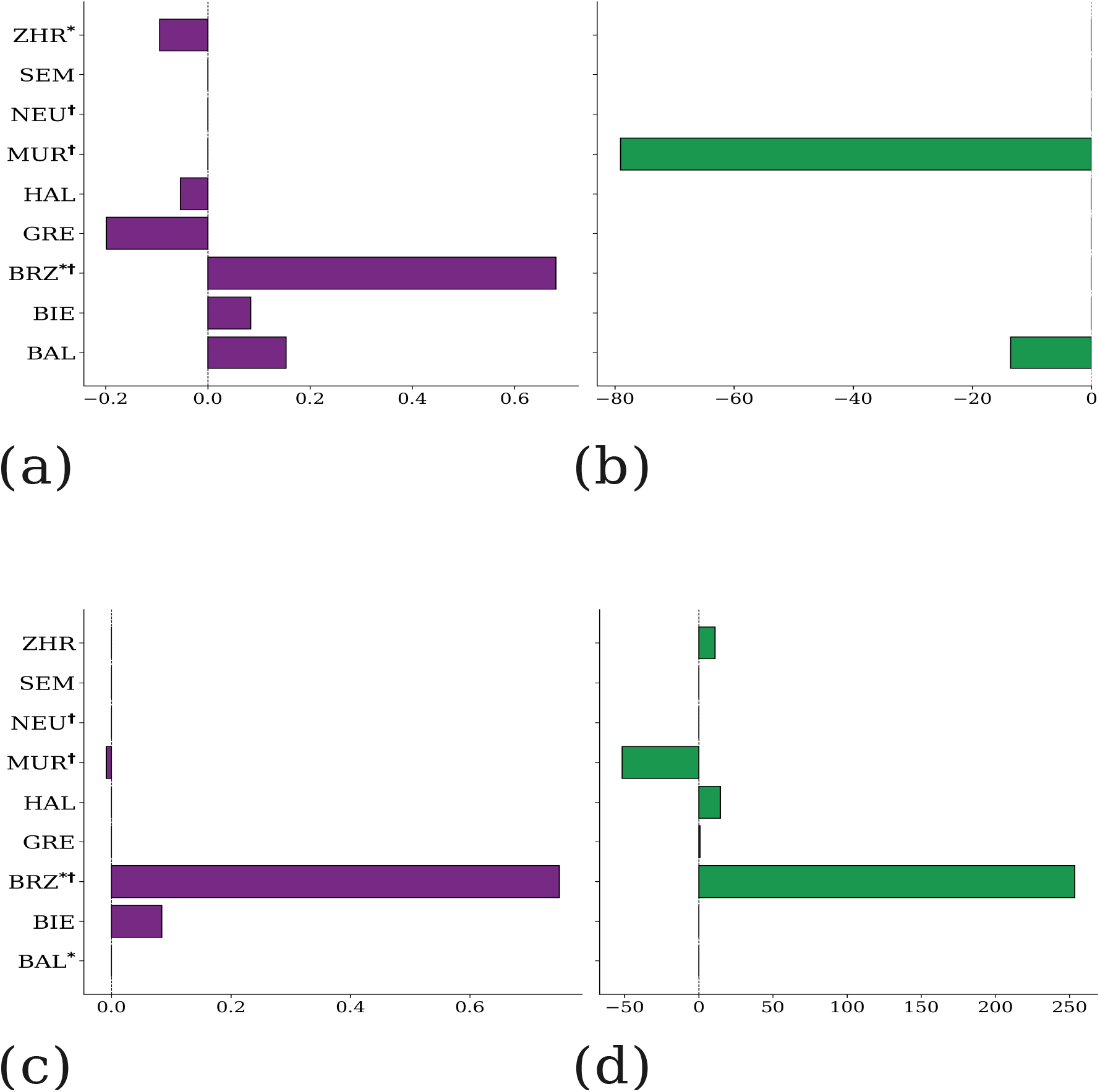
SARIMAX coefficients for asymptotic stability under orthogonalized environmental drivers. Panels report lake-specific coefficients for log(−*α*) (with *α* the real part of the dominant eigenvalue) estimated from SARIMAX models using lagged temperature and phosphate covariates. Top row: models including temperature lags and phosphate residualized on the temperature lag block 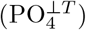. Bottom row: models including phosphate lags and temperature residualized on the phosphate lag block 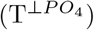. Left column (a,c) shows coefficients for the temperature predictor (temperature in the top row; 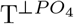in the bottom row), and right column (b,d) shows coefficients for the phosphate predictor 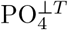 in the top row; phosphate in the bottom row). For each lake, bars summarize the average coefficient across lags that show the strongest statistical support in that lake. Lake acronyms denote: BAL (Lake Baldegg), BIE (Lake Biel), BRZ (Lake Brienz), GRE (Lake Greifen), HAL (Lake Hallwil), MUR (Lake Murten), NEU (Lake Neuenburg), SEM (Lake Sempach), and ZHR (Lake Zurich). An asterisk (*) denotes lakes where residual diagnostics failed, and a dagger () denotes lakes where the absolute PO_4_ change is < 5% of the maximum observed variation.

**S8 Fig.**
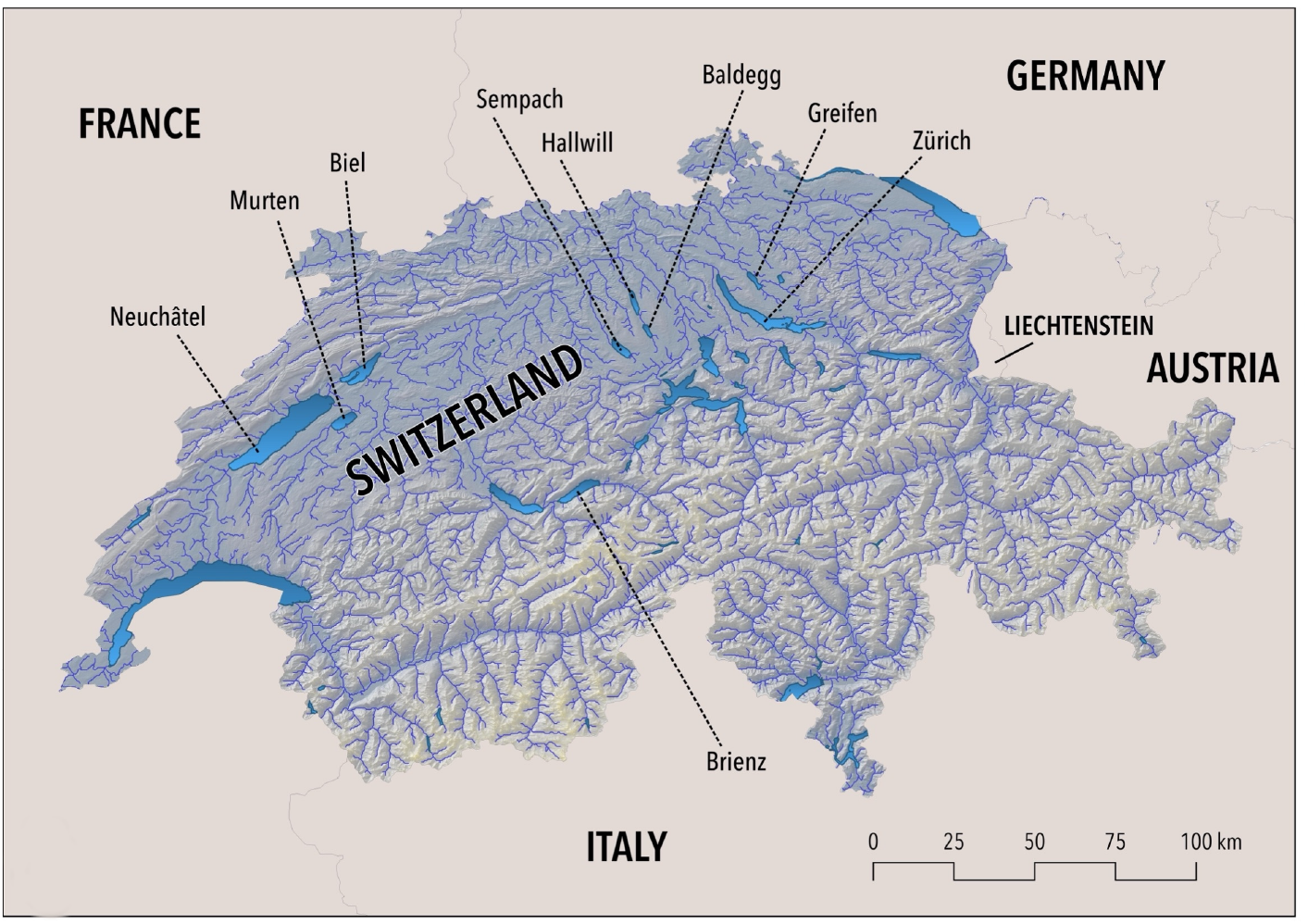
Location of the studied lakes across Switzerland, highlighting the major basins included in the analysis.

**S9 Fig.**
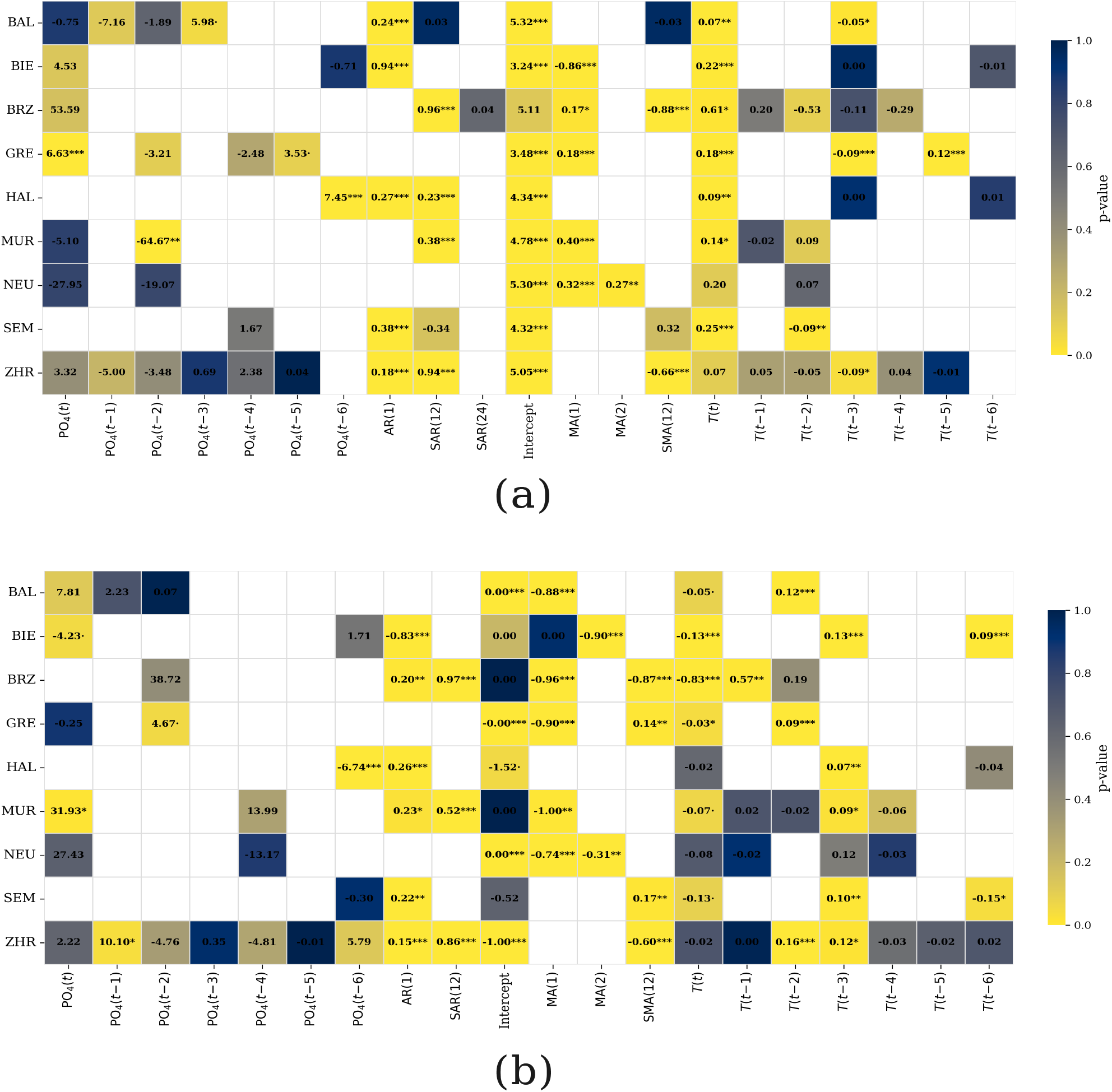
Estimated regression coefficients from lake-specific SARIMAX models with orthogonalized phosphate. Rows correspond to lakes and columns to model terms, including temperature covariates at multiple lags, autoregressive and moving-average components, and phosphate residualized with respect to the temperature lag block (i.e., 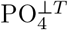 ; see Methods). Numbers report coefficient estimates, with different type I error estimates indicated by asterisks (^*\**^*p* < 0.05, ^*\*\**^*p* < 0.01, ^*\*\*\**^*p* < 0.001; ·*p* < 0.1). Cell color encodes the associated *p*-value on a continuous scale from 0 to 1, with darker colors indicating stronger statistical support. Panel (a) reports coefficients for reactivity, whereas panel (b) reports coefficients for *t*_max_. Blank cells indicate terms not retained in the selected model for a given lake. Lake acronyms denote: BAL (Lake Baldegg), BIE (Lake Biel), BRZ (Lake Brienz), GRE (Lake Greifen), HAL (Lake Hallwil), MUR (Lake Murten), NEU (Lake Neuenburg), SEM (Lake Sempach), and ZHR (Lake Zurich).

**S10 Fig.**
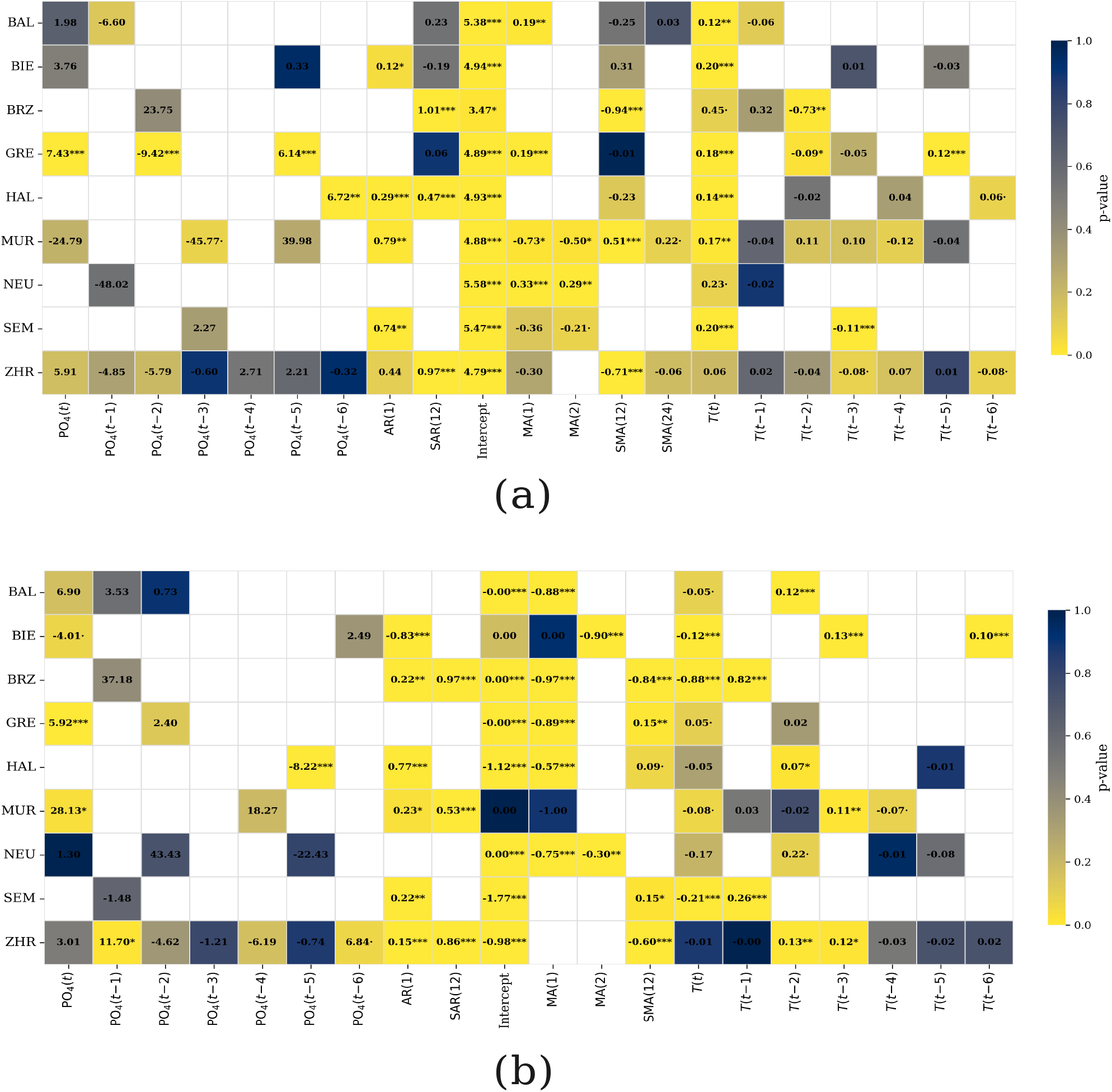
Estimated regression coefficients from lake-specific SARIMAX models with orthogonalized temperature. Rows correspond to lakes and columns to model terms, including phosphate covariates at multiple lags, autoregressive and moving-average components, and temperature residualized with respect to the phosphate lag block (i.e., 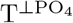 ; see Methods). Numbers report coefficient estimates, with different type I error estimates indicated by asterisks (^*\**^*p* < 0.05, ^*\*\**^*p* < 0.01, ^*\*\*\**^*p* < 0.001; ·*p* < 0.1). Cell color encodes the associated *p*-value on a continuous scale from 0 to 1, with darker colors indicating stronger statistical support. Panel (a) reports coefficients for reactivity, whereas panel (b) reports coefficients for *t*_max_. Blank cells indicate terms not retained in the selected model for a given lake. Lake acronyms denote: BAL (Lake Baldegg), BIE (Lake Biel), BRZ (Lake Brienz), GRE (Lake Greifen), HAL (Lake Hallwil), MUR (Lake Murten), NEU (Lake Neuenburg), SEM (Lake Sempach), and ZHR (Lake Zurich).

**S11 Fig.**
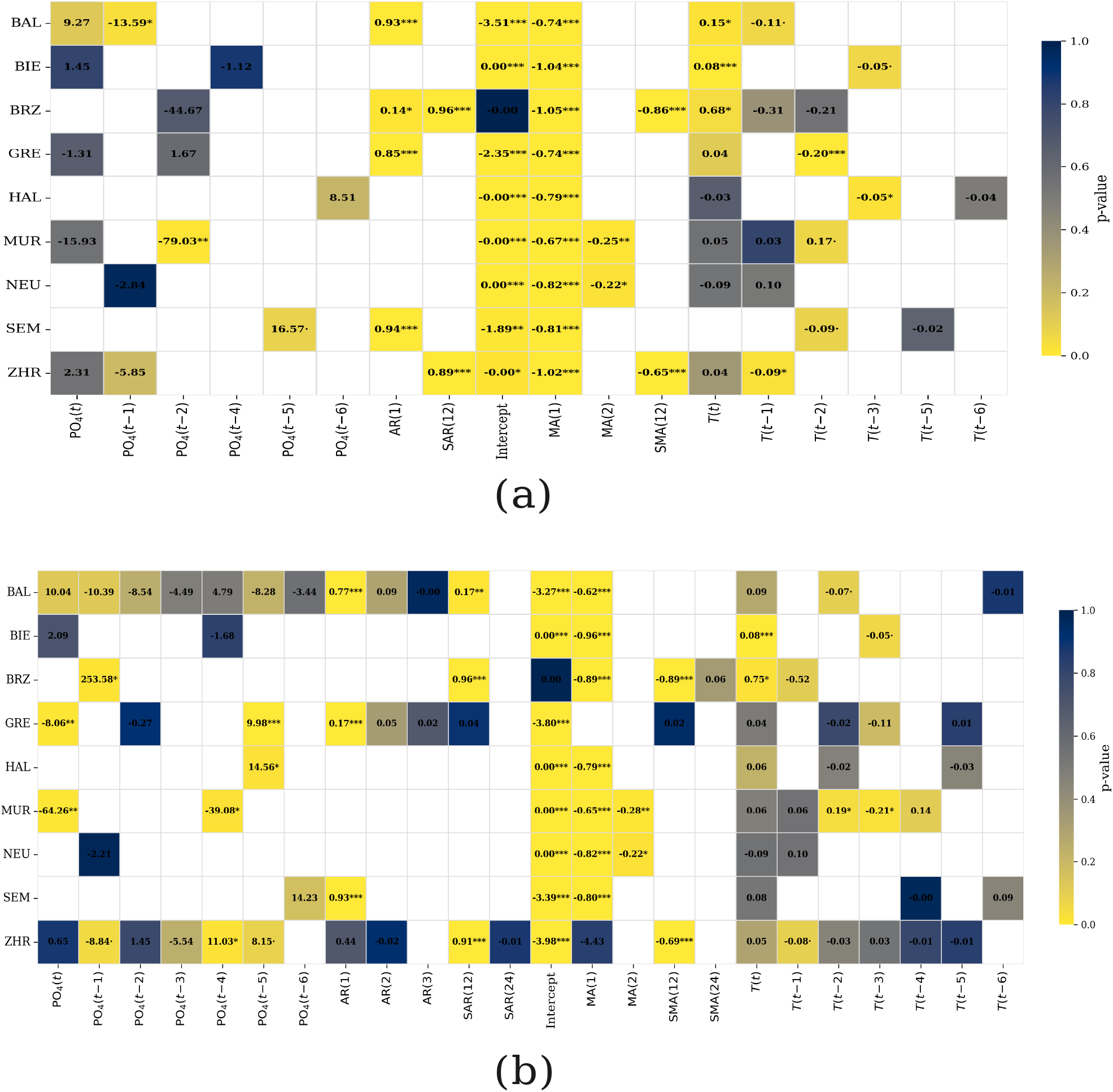
Estimated regression coefficients from lake-specific SARIMAX models for asymptotic stability. Rows correspond to lakes and columns to model terms, including lagged temperature and phosphate covariates, as well as autoregressive and moving-average components. Coefficients refer to log(−*α*), where *α* is the real part of the dominant eigenvalue. Panel (a) reports results from models including temperature lags and phosphate residualized with respect to the temperature lag block 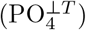, whereas panel (b) reports results from models including phosphate lags and temperature residualized with respect to the phosphate lag block 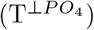. Numbers indicate coefficient estimates, with different type I error estimates denoted by asterisks (^*\**^*p* < 0.05, ^*\*\**^*p* < 0.01, ^*\*\*\**^*p* < 0.001; · *p* < 0.1). Cell color encodes the associated *p*-value on a continuous scale from 0 to 1, with darker colors indicating stronger statistical support. Blank cells indicate terms not retained in the selected model for a given lake. Lake acronyms denote: BAL (Lake Baldegg), BIE (Lake Biel), BRZ (Lake Brienz), GRE (Lake Greifen), HAL (Lake Hallwil), MUR (Lake Murten), NEU (Lake Neuenburg), SEM (Lake Sempach), and ZHR (Lake Zurich).

### SI-G. Sensitivity analysis: temperature-scaled bioenergetic parameters

The reactivity and time-to-peak-response time series reported in the main text were computed with bioenergetic parameters fixed to the values of Boit et al. [36]. A natural concern is that warming should, in reality, modify those parameters according to known thermal scaling [3, 4, 13]. If the fixed-parameter result reflects only the temperature dependence of the observed biomass distribution, and not also the temperature dependence of the metabolic and ingestion rates that connect biomasses to interaction strengths, then the central finding could be parameterization-specific.

To probe this concern, we re-ran the entire pipeline with the bioenergetic parameters made explicitly temperature-dependent through Boltzmann-Arrhenius scaling, and we compared the resulting reactivity time series to the fixed-parameter version lake by lake.

We treat the Boit et al. [36] values as reference rates at *T*_ref_ = 20 °C (*T*_ref,K_ = 293.15 K) and apply the correction

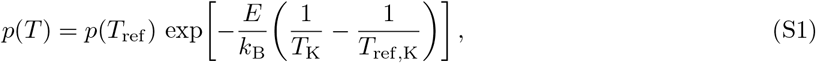

with *k*_B_ = 8.617 × 10^*−*5^ eV K^*−*1^.

#### Parameters scaled

- Producer intrinsic growth rates *r*_*i*_ (activation energy *E*_*r*_ = 0.47 eV) [3, 13];
- Consumer metabolic rates *x*_*i*_ (activation energy *E*_*x*_ = 0.65 eV), standard for ectotherm metabolism [4, 13];
- Maximum-consumption multiplier *y* (effective activation energy *E*_*y*_ = *E*_*J*_ − *E*_*x*_ = 0.10 eV, with *E*_*J*_ = 0.75 eV for ingestion) was scaled in a separate “mechanistic” run; the main comparison reported below uses the conservative version with *y* fixed.

Assimilation efficiencies *e*_*i*_, prey preferences *ω*_*ij*_ (i.e. topology), carrying capacity *k* and half-saturation constant *B*_0_, and self-limitation *m* were not rescaled, both because their thermal dependence is more variable across taxa and because they would otherwise compound several scaling assumptions in a single sensitivity analysis. Closure coefficients *α*_*di*_ and *α*_*id*_ were recomputed after scaling, because they are derived from the mass balance of biological fluxes; copying the unscaled values would violate mass balance under the new rates. For each lake-month we (i) loaded observed biomasses ***B***(*t*) and water temperature *T* (*t*), (ii) rescaled *r*_*i*_ and *x*_*i*_ via Eq. (S1), (iii) computed biological fluxes with the rescaled parameters, (iv) recomputed mass-balance closure *α*_*di*_, *α*_*id*_, (v) constructed the Jacobian ***J*** _thermal_(*t*), and (vi) extracted *ρ*_thermal_(*t*). Where lake temperature was missing for a given month, the pipeline fell back to unscaled reference parameters; this affected only a small number of observations.

#### Comparison with the fixed-parameter

We matched the fixed-parameter and thermal-parameter outputs by (lake, year, month), obtaining 2,949 paired lake-months across the nine lakes for which the comparison could be carried out.

Across all 2,949 matched lake-months, the rank correlation between fixed-parameter and thermal-parameter reactivity is Spearman *ρ* = 0.988 (*p* < 10^*−*3^), and the linear correlation is Pearson *r* = 0.971 (*p* < 10^*−*3^).

Table S5 reports the same comparison lake by lake. Spearman *ρ >* 0.94 in every lake, indicating that the rank ordering of monthly reactivity is essentially preserved when bioenergetic parameters are made temperature-dependent. Pearson *r* is also high but lower for a few lakes (BAL, HAL, NEU), reflecting amplitude differences rather than ordering disagreements.

Absolute reactivity is systematically lower under thermal scaling (mean relative change ≈−57% globally).

This is the expected direction: most lake temperatures lie below the reference *T*_ref_ = 20 °C, which drives down the rescaled metabolic rates (*x*_*i*_) and growth rates (*r*_*i*_), and therefore the magnitude of the off-diagonal Jacobian entries. The rank ordering of monthly reactivity within each lake, and the cross-lake pattern of warming-driven reactivity changes, are preserved (Fig. S12).

**S5 Table.**
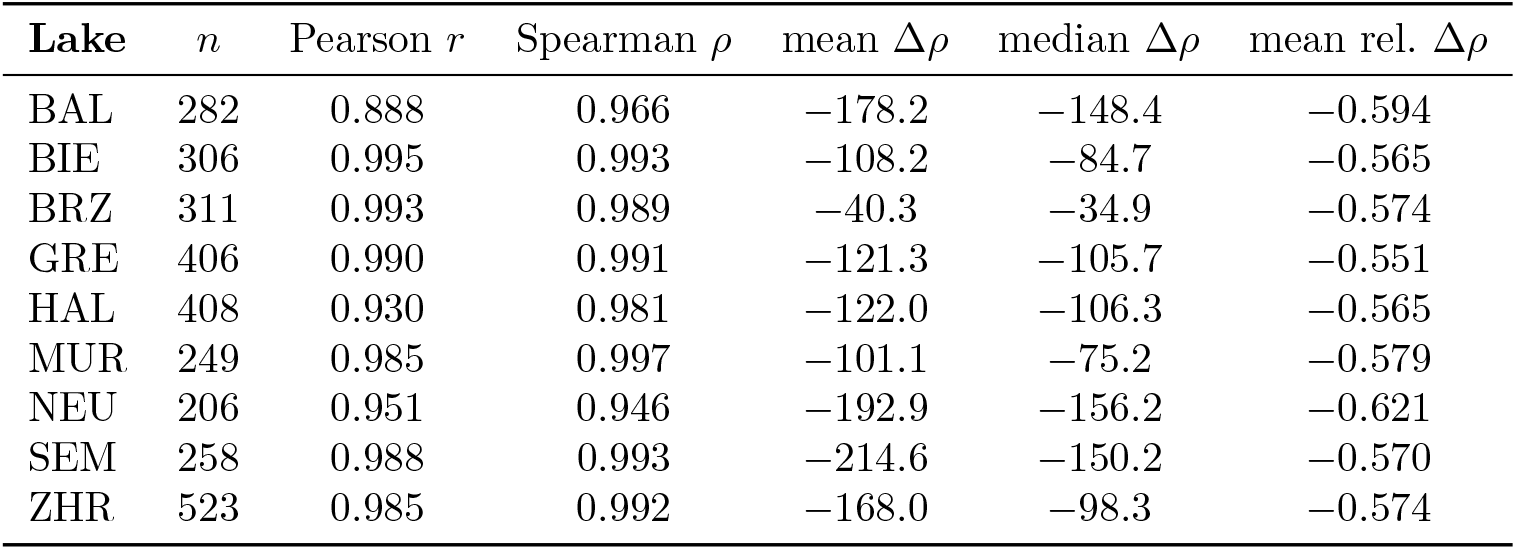
Lake-by-lake correlation between fixed-parameter and thermal-parameter reactivity time series. *n* is the number of matched lake-months; Δ*ρ* = *ρ*_thermal_ −*ρ*_fixed_; relative change is Δ*ρ/ρ*_fixed_. Spearman *ρ* is the rank correlation.

The qualitative warming-reactivity association reported in the main text is not an artifact of holding bioenergetic parameters fixed. Allowing those parameters to scale with temperature according to standard thermal-ecology activation energies preserves both the within-lake temporal structure and the cross-lake pattern of reactivity change. We interpret this as an internal mechanistic check on the central result.

**S12 Fig.**
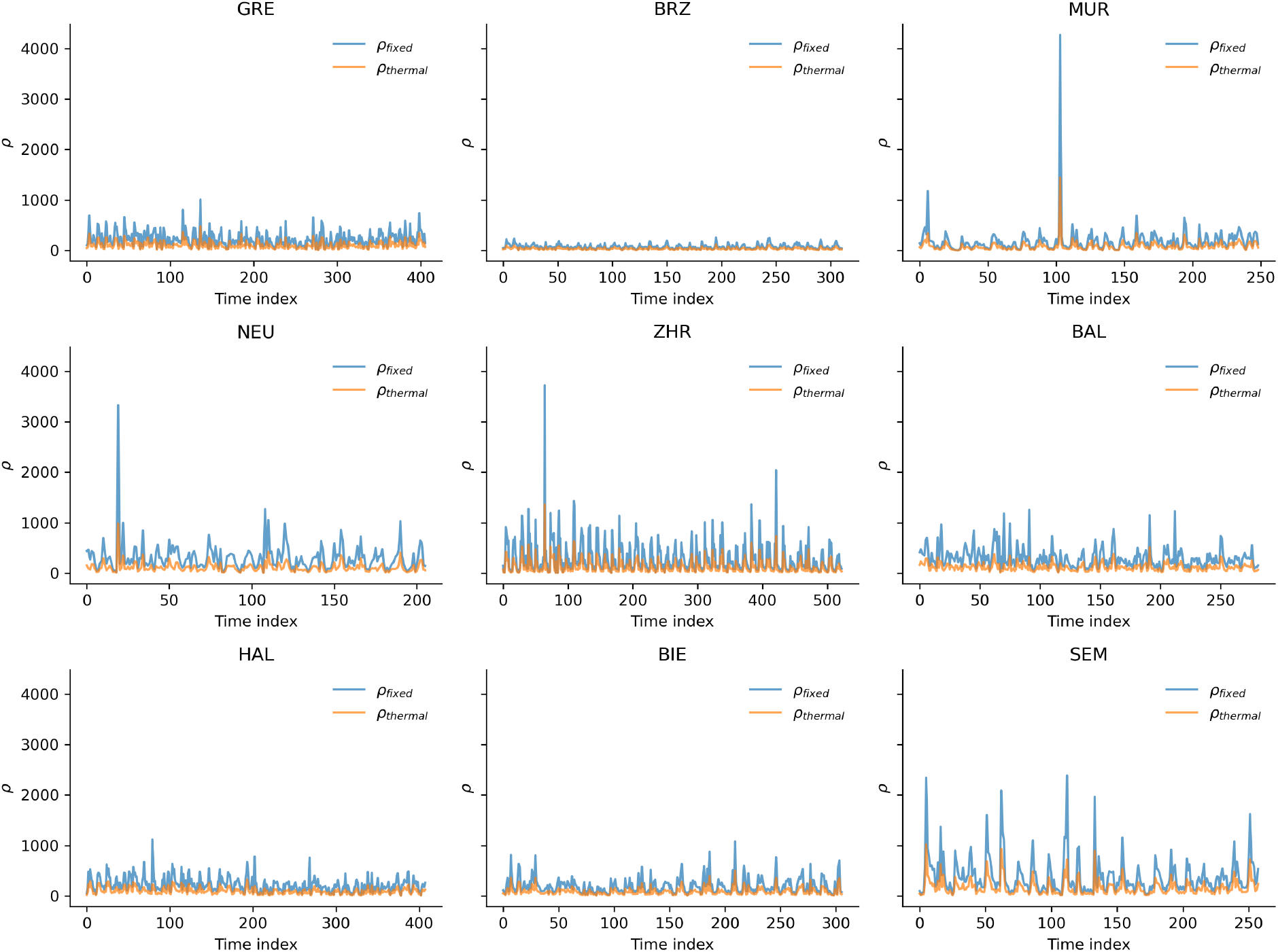
Fixed-parameter (*ρ*_fixed_) versus temperature-scaled (*ρ*_thermal_) reactivity, lake by lake. Each panel shows the monthly time series of fixed-parameter reactivity (blue) and thermal-parameter reactivity (orange) across the matched lake-months. Absolute values of *ρ*_thermal_ are systematically lower because most lake temperatures sit below the reference *T*_ref_ = 20 °C, but the rank ordering and within-lake temporal structure are preserved (Spearman *ρ >* 0.94 in every lake; Table S5). The qualitative warming-reactivity association reported in the main text is therefore not an artifact of holding bioenergetic parameters fixed.

## Notes

### Competing Interest Statement

The authors have declared no competing interest.

